# Discovery of SARS-CoV-2 papain-like protease (PL^pro^) inhibitors with efficacy in a murine infection model

**DOI:** 10.1101/2024.01.26.577395

**Authors:** Michelle R. Garnsey, Matthew C. Robinson, Luong T. Nguyen, Rhonda Cardin, Joseph Tillotson, Ellene Mashalidis, Aijia Yu, Lisa Aschenbrenner, Amanda Balesano, Amin Behzadi, Britton Boras, Jeanne S. Chang, Heather Eng, Andrew Ephron, Tim Foley, Kristen K. Ford, James M. Frick, Scott Gibson, Li Hao, Brett Hurst, Amit S. Kalgutkar, Magdalena Korczynska, Zsofia Lengyel-Zhand, Liping Gao, Hannah R. Meredith, Nandini C. Patel, Jana Polivkova, Devendra Rai, Colin R. Rose, Hussin Rothan, Sylvie K. Sakata, Thomas R. Vargo, Wenying Qi, Huixian Wu, Yiping Liu, Irina Yurgelonis, Jinzhi Zhang, Yuao Zhu, Lei Zhang, Alpha A. Lee

## Abstract

Vaccines and first-generation antiviral therapeutics have provided important protection against coronavirus disease 2019 (COVID-19) caused by severe acute respiratory syndrome coronavirus 2 (SARS-CoV-2). However, there remains a need for additional therapeutic options that provide enhanced efficacy and protection against potential viral resistance. The SARS-CoV-2 papain-like protease (PL^pro^) is one of two essential cysteine proteases involved in viral replication. While inhibitors of the SARS-CoV-2 main protease (M^pro^) have demonstrated clinical efficacy, known PL^pro^ inhibitors have to date lacked the inhibitory potency and requisite pharmacokinetics to demonstrate that targeting PL^pro^ translates to *in vivo* efficacy in a preclinical setting. Herein, we report the machine learning-driven discovery of potent, selective, and orally available SARS-CoV-2 PL^pro^ inhibitors, with lead compound PF-07957472 (**4**) providing robust efficacy in a mouse-adapted model of COVID-19 infection.

## Introduction

Coronaviruses are a group of positive-sense single-stranded RNA viruses known to cause human pathologies, ranging from mild to moderate upper-respiratory tract illnesses (229E, NL63, OC43, and HKU1) to global outbreaks (SARS-CoV-1, Middle East Respiratory Syndrome-CoV, and SARS-CoV-2) [1,2]. In particular, the COVID-19 pandemic caused by SARS-CoV-2 has led to over 14 million estimated excess deaths worldwide as of late 2023. Highly effective vaccines have provided community-wide protection, while oral antivirals are currently approved for adults with mild-to-moderate COVID-19 who are at high risk for progression to severe disease.

Currently, SARS-CoV-2 antivirals, which have been clinically approved or are in late-stage development, are directed against either the RNA-dependent polymerase (RdRp) or the main protease (M^pro^) [3]. With the possibility of resistance developing over time [4–6], there is a need for additional oral antivirals directed against currently undrugged viral targets.

The SARS-CoV-2 papain-like protease (PL^pro^) is a cysteine protease and a domain in the coronavirus non-structural protein 3 (Nsp3). Conserved across coronaviruses, PL^pro^, along with M^pro^, is responsible for polyprotein processing, which is required for the generation of a functional viral replicase [7]. Additionally, PL^pro^ functions as a deubiquitinase/deISGylase and is thought to modulate host innate immune pathways via cleavage of the post translational modifications ubiquitin and ISG15 from host proteins as an evasion mechanism [8]. As such, PL^pro^ is hypothesized to be an essential viral enzyme [9], and PL^pro^-targeting antivirals may offer a differentiated mechanism of action compared to antivirals in the current therapeutic landscape.

However, there are two key missing elements in understanding the therapeutic potential of targeting PL^pro^: 1) whether inhibition of PL^pro^ can result in cellular antiviral activity at therapeutically meaningful potency and 2) whether potent PL^pro^ inhibition can translate to *in vivo* efficacy with compounds that have a well-characterized selectivity profile and favourable pharmacokinetic attributes in preclinical species. In particular, animal disease models of COVID-19 can help establish the translation between *in vitro* PL^pro^ inhibition and *in vivo* viral suppression, as well as potentially shed light on the interplay between infection and immune response [10]. However, identification of PL^pro^ inhibitors suitable for *in vivo* validation is challenging with respect to achieving the desired combination of the requisite PL^pro^ potency, cellular antiviral activity, and appropriate preclinical pharmacokinetics as maintaining trough plasma concentrations in vivo at multiples of the cellular antiviral potency is necessary [11,12]. Thus far, compounds in the literature include compounds developed for SARS-CoV-1 PL^pro^ [9,13–16], which also show activity against SARS-CoV-2 PL^pro^ (most notably GRL0617, Compound **1**), as well as compounds identified more recently to combat the SARS-CoV-2 pandemic. While valuable starting points, these molecules have shown only weak cellular antiviral potencies in the low-to-high micromolar range with minimal information on *in vitro* (or *in vivo*) disposition characteristics in animals and/or human reagents.

Herein, we report the machine learning-driven discovery of potent, selective, and orally bioavailable SARS-CoV-2 PL^pro^ inhibitors with robust efficacy in a mouse-adapted model of COVID-19 infection.

### Discovery of potent PL^pro^ inhibitors

Given the expected need for rapid improvements in relevant physicochemical properties during the rapidly evolving pandemic, a key approach of our medicinal chemistry strategy was data-driven and algorithmic. AI and machine learning (ML) have seen increased use in small molecule drug discovery in recent years, with the advent of generative chemistry and multiparameter optimization approaches benefitting from the broader advances and resurgence in ML technology [17,18]. As part of our machine learning driven approach, we used parallel high throughput chemistry to rapidly scan large regions of chemical space, probe multiple synthetically accessible vectors, and diversely select compounds for each library based on bioactivity predictions [19,20]. Critical to the success of the collaboration between Pfizer and PostEra to ensure efficiency was to leverage the integration of CRO networks as described in Figure S1. The workflow set up allowed for speed in execution and delivery of compounds and data to drive SAR.

We started from known compounds such as the widely studied GRL0617 (**1**), a non-covalent inhibitor active against SARS-CoV-1, subsequently found to be active against SARS-CoV-2 PL^pro^ [13,14] (**Figure 1A**). Compound **1** inhibits recombinant SARS-CoV-2 PL^pro^ with a 857 nM inhibition constant K_i_ in our fluorescence resonance energy transfer (FRET)–based substrate cleavage assay. In a cellular context, **1** showed weak antiviral effects (half-maximal effective concentration (EC_50_) of 51.9 μM) measured by monitoring the cytopathic effect (CPE) in Vero E6 cells in the presence of the P-glycoprotein efflux inhibitor CP-100356 [12]. To improve upon the existing chemical matter, we identified two regions for optimization: the naphthalene ring system and the aniline substituent. Our first round of medicinal chemistry efforts focused on elaboration of the naphthalene ring. Scanning available building blocks led to the discovery of a quinoline substituent as a suitable replacement. A large Suzuki coupling library was designed using machine learning to explore the 2-position (**Figure 1B**) while balancing predicted bioactivity, molecular diversity, and the synthetic requirements of building blocks for a successful execution of the parallel library chemistry approach. Our library led to the identification of the *N*-methyl pyrazole (**2**) derivative with an order-of-magnitude improvement in PL^pro^ K_i_, even in the absence of the aniline functional group.

**Figure 1:**
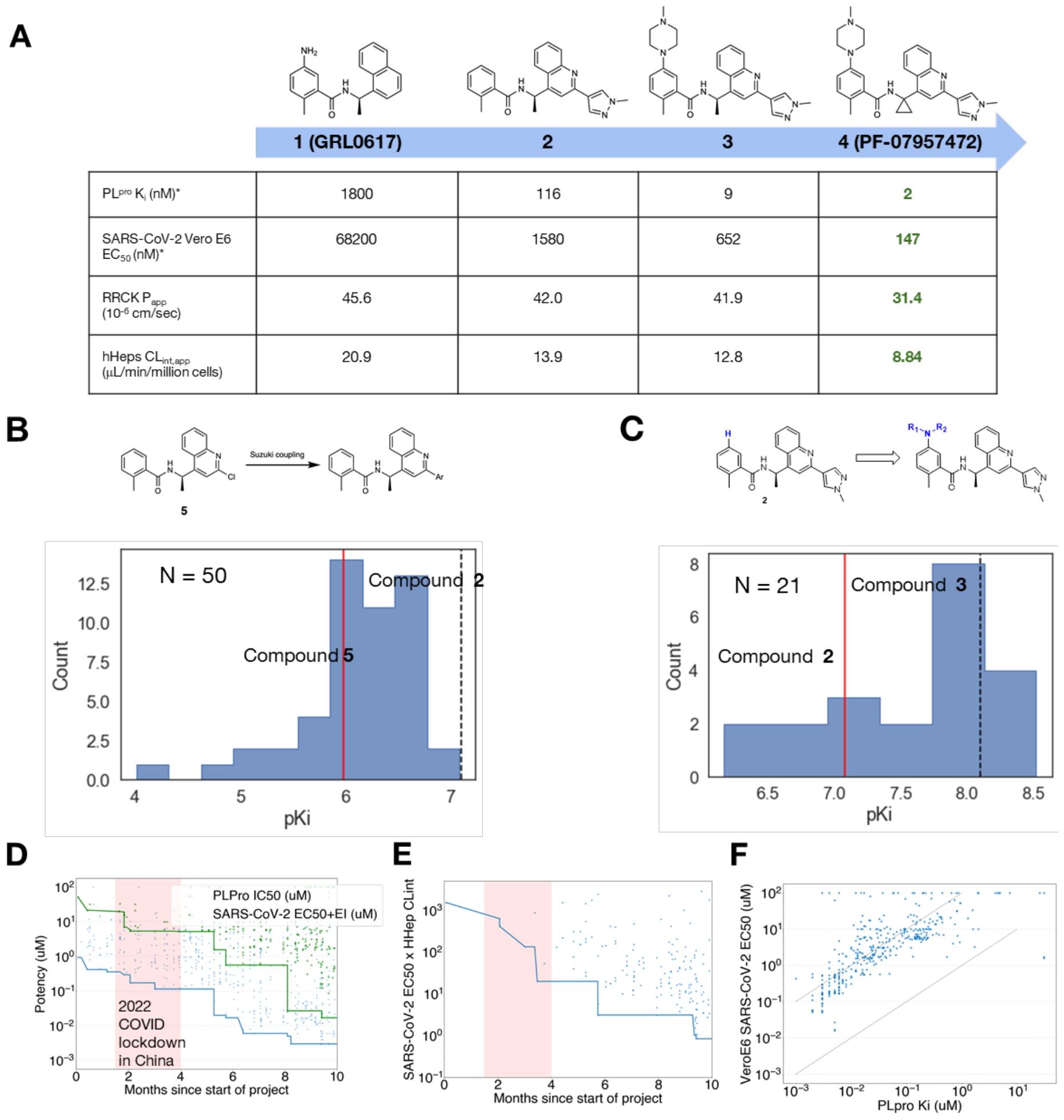
Discovery of PLpro inhibitors guided by machine learning. (A) Summary of key compounds in our discovery campaign with their biochemical potency, cell antiviral potency in Vero E6 cells containing the P-glycoprotein inhibitor CP-100356 (2 μM), passive permeability determined by Ralph Russ canine kidney cells (RRCK) [23] and apparent intrinsic clearance (CLint,app) obtained from metabolic stability studies in cryopreserved human hepatocytes [24]. *all values provided are the geometric mean of at least three replicates. (B)-(C) show the potency of compounds made in the Suzuki and C-N coupling libraries. The red line denotes the potency of the compound that inspired the library. (D) Biochemical (blue) and cell (green) potency over time, and (E) product between potency and CLint,app in human hepatocytes [24] of synthesized compounds plotted over time. In (D) and (F), each datapoint is a compound where we track the date of synthesis completion, and the solid lines show the running minimum. As our synthetic chemistry efforts were based in China, COVID lockdown there precipitated a marked disruption to campaign progress. (F) PLpro inhibition translates to antiviral activity. Ki and inhibition are correlated on a cellular level (in presence of 2 μM of Pgp inhibitor CP-100356 to counteract the high levels of Pgp efflux in Vero E6 cells [12]). The dotted lines show EC50 = Ki and 100x Ki respectively.

Our second round of optimization focused on modifying the aniline motif. To this end, we constructed libraries based on C-N coupling (**Figure 1C**), which led to the identification of methyl piperazine substituent with another order-of-magnitude improvement in K_i_, while also providing a 2-fold improvement in viral CPE potency in the Vero E6 assay. Finally, rigidification of the scaffold via introduction of a geminal-cyclopropyl group in lieu of the pendant methyl substituent, resulted in the lead compound (PF-07957472, **4**) with an additional 3-fold improvement in biochemical and cellular antiviral potencies, and also a reduction in the apparent intrinsic clearance (CL_int,app_) in a metabolic stability assay using human hepatocytes [21].

The data-driven strategy led to a compressed timeline (**Figure 1D**) wherein month-on-month improvements in biochemical potency against PL^pro^ and cellular antiviral activity were achieved. With a desire to establish proof-of-mechanism in an animal model of COVID-19 and potentially identify chemical lead matter with clinical candidate-like qualities, a crucial indicator of campaign progress was the concomitant optimization of potency and metabolic stability of new lead compounds. This balance could be captured by evaluating the product between cellular antiviral activity (EC_50_) and metabolic CL_int,app_ estimated in human hepatocyte incubations, a quantitative score inspired by a fit-for-purpose human dose prediction (see SI section Methodology for Multi-Parameter Optimization Scoring) [22]. The quantitative score revealed a steady reduction over time (**Figure 1E**), showing that desired PL^pro^ inhibitory potency and cellular antiviral activity could be achieved for compounds with reasonably low CL_int,app_ values in human hepatocytes. Additional confidence in PL^pro^ as an antiviral target was evident from the strong correlation between enzymatic inhibition of PL^pro^ and cellular antiviral activity (**Figure 1F**). In general, the cellular antiviral EC_50_ was observed to be approximately 2 orders of magnitude weaker than the corresponding biochemical K_i_ against recombinant PL^pro^, which is not unexpected given biochemical-to-cell potency translation observed on similar protease targets for reversible inhibitors [12], though the precise reasons for this significant drop off remain speculative.

### PF-07957472 is a suitable *in vivo* tool compound

A crystal structure of SARS-CoV-2 PL^pro^ in complex with **4** was determined to 2.59 Å (**Table S2**). This structure reveals that **4** engages a region on PL^pro^ that overlaps with the substrate binding site but does not extend to the catalytic triad active site (**Figure S2A-B**). Compound **4**, like GRL0617, binds in the pocket formed by the flexible BL2 loop and forms a critical hydrogen bond with Asp164 [14,25] (**Figure 2A**). The binding pose assumed by **4** differs from that of GRL0617 in the following ways: (1) the *N*-methyl pyrazole-quinoline aromatic system in **4**, in place of the naphthalene moiety, gains greater surface area coverage of the hydrophobic ‘shelf’, formed by Pro247 and Pro248, and more efficient T-shaped π-stacking interactions with Tyr268 from a ∼30° tilt at the quinoline core towards this residue (**Figure S2C**); (2) the protonated piperazinyl *N*-methyl amine engages Glu167 in a salt bridge (3.6 Å), while the analogous primary amine of the aniline of GRL0617 is 6 Å away from Glu167 (**Figure S2D**); (3) The cyclopropyl is a larger hydrophobic substitution that better engages Tyr264 via CH-π interactions, while its hydrogen atoms, polarized due to conformational strain, can also interact with the polar residues in the pocket such as Thr301 and Asp164, a feature that is missing in GRL0617, which has a smaller methyl group at this position (**Figure S2E**). These optimized protein-ligand interactions rationalize the significantly improved potency observed for **4**, compared to GRL0617 (**Figure 2A**; **Figure S2C-E**).

**Figure 2:**
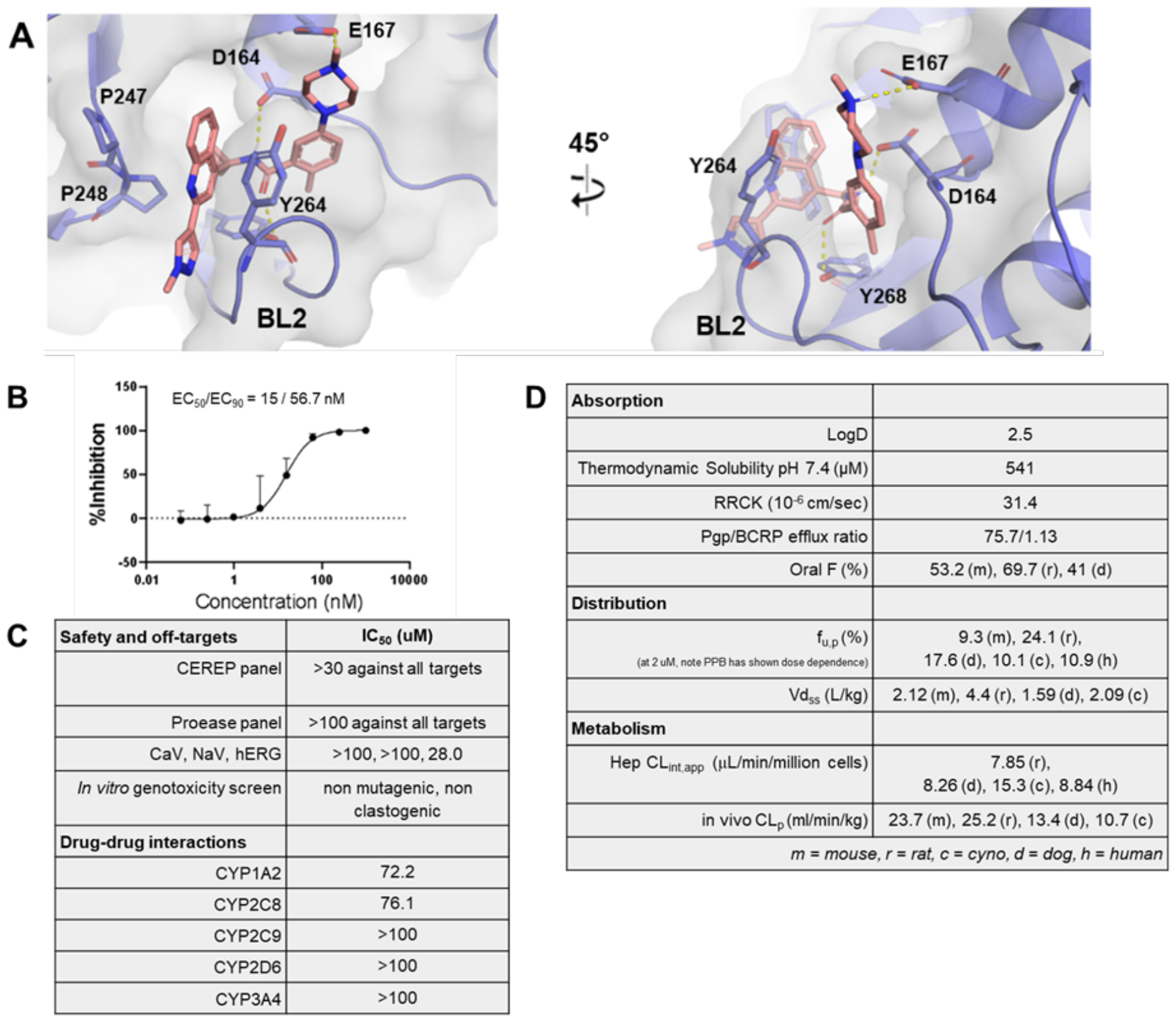
Compound 4 is an antiviral lead and a suitable in vivo tool compound. (A) X-ray crystal structure of **4** (pink) bound to SARS-CoV-2 PL^pro^ (blue, grey surface representation), highlighting the critical protein-ligand interactions key to high affinity. (B) Dose-response curve of **4** assayed in a human airway epithelial cell model of SARS-CoV-2 infection [12]. The fraction unbound of **4** in the dNHBE media was measured to be 0.916, assumed to be 1 for modeling purposes.(C) Profiling **4** through safety panels and CYP panels reveal no clear off-target liabilities. (D) In vitro and in vivo absorption, distribution and metabolism properties across multiple species.

We profiled **4** in a primary human airway epithelial (dNHBE) cell infection model, a physiologically relevant system that had shown better *in vitro-in vivo* correlation than Vero E6 cells to the observed efficacy with published Mpro inhibitors [26]. Compound **4** demonstrated potent viral CPE in dNHBE cells (EC_50_ = 15 nM) (**Figure 2B, Figure S4**). To identify potential off-target activity, we profiled **4** through standard safety pharmacology panels as well as comprehensive protease panels (further details in the supplementary info). We detected no major off-target activity, with only moderate human ether-a-go-go (hERG) potassium ion channel (product of the hERG gene; Kv11.1) activity (IC_50_ = 28.0 μM) which was considered marginal, given the potent antiviral activity in cells. Compound **4** was devoid of potent reversible inhibition of major cytochrome P450 (CYP) isoforms, including the major human constitutive CYP enzyme CYP3A4 (**Figure 2C**). The observation that selective PL^pro^ active site inhibition leads to low nM antiviral activity in a primary cellular system provides additional evidence for the essentiality of PL^pro^ in viral replication.

Compound **4** was shown to have favorable *in vitro* ADME properties including high thermodynamic solubility and passive permeability in the RRCK assay, and a low metabolic CL_int,app_ in human hepatocytes (**Figure 2D**). Following intravenous administration to preclinical species (mice, rats, dog, and monkeys), **4** demonstrated moderate plasma clearances (CL_p_) across species and high steady state distribution volumes (Vd_ss_). Despite being a Pgp substrate [27], **4** demonstrated moderate oral bioavailability (F) and a high fraction of the oral dose absorbed across the preclinical species studied, in a relatively straightforward formulation comprising of 0.5% methylcellulose in water containing 2% Tween® 80. The oral pharmacokinetics of **4** were encouraging, particularly against the backdrop of established pharmacokinetic-pharmacodynamic relationships for SARS-CoV-2 protease inhibitors (and protease inhibitors for HIV and HCV as well [11]), which require trough or minimum plasma concentrations (C_min_) to be maintained above cellular EC_90_ during the entire treatment duration [3].

### PL^pro^ inhibitor PF-07957472 is efficacious in a murine SARS-CoV-2 infection model

Given its promising antiviral activity and acceptable rodent oral F, a multi-dose mouse pharmacokinetics study was conducted with orally administered **4** (30, 150, and 500 mg/kg), which demonstrated that unbound systemic exposures were considerably higher than the dNHBE EC_90_ of 56.7 nM. Moreover, **4** was tolerated at the highest dose studied.

Based on the findings from the preliminary dose-range finding studies, antiviral activity of **4** was examined in a mouse-adapted SARS-CoV-2 (SARS-CoV-2 MA10) model [28], following twice daily (BID) oral administration at 20, 50, and 150 mg/kg. Nirmatrelvir was included as a positive control and dosed orally at 1000 mg/kg (BID), which had proven to be efficacious in this mouse model [12].

In the efficacy study, mice were infected four hours prior to the first dose, dosed BID for four days and then viral lung titers were measured. Compound **4** caused statistically significant reduction in lung viral titers at four days post infection for both the 50 and 150 mg/kg dose groups at unbound systemic exposures (C_min_) that maintained or exceeded the dNHBE EC_90_ through the dosing period (**Figure 3A, B and D**). The observed efficacy is comparable to that observed with the SARS-CoV-2 main protease inhibitor Nirmatrelvir (**Figure 3B and D**). While infected and untreated mice had robust infection at Day 4, half of the mice in the 150 mg/kg dose had viral levels reduced to the limit of viral detection. Additionally, BID treatment with **4** protected mice from weight loss compared with the infected mice in the vehicle group, which showed ∼10% body weight loss following infection as expected (**Figure 3C**). Overall, these findings confirmed that PL^pro^ inhibition is indeed effective at reducing SARS-CoV-2 viral replication in mouse lungs, and that the PL^pro^ inhibitor **4** is an effective *in vitro* and *in vivo* tool compound for further studies of this novel mechanism of antiviral effect.

**Figure 3:**
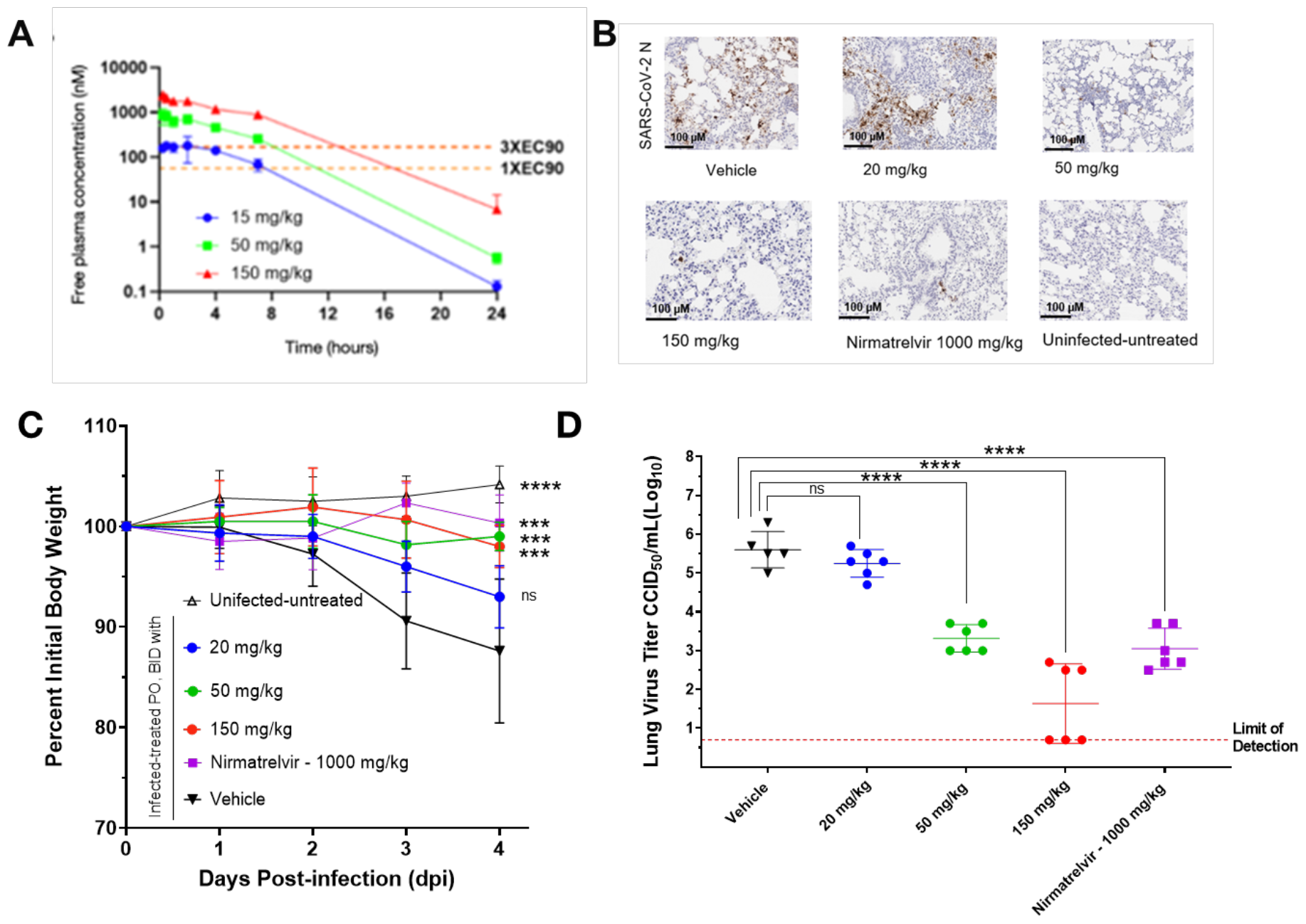
PLpro inhibitor 4 is efficacious in a murine SARS-CoV-2 infection model. (A) Dose-Range Finding studies showed that the selected doses cover a range of trough concentrations from sub-therapeutic to therapeutic levels. (B) Representative IHC images from lung histopathology of mouse-adapted SARS-CoV-2 infection 4 days post dose of **4** at 20, 50, and 150 mg/kg, Nirmatrelvir at 1000 mg/kg, vehicle, or uninfected, untreated animals. (C) Compound **4** protected mice from weight loss, with roughly 10% weight loss seen in the vehicle arm consistent with data in [12]. (D) Compound **4** led to a statistically significant and dose-dependent reduction in Day 4 lung viral titers. Titers were plotted as mean log10 CCID_50_/ml ± SEM, with data analysis and significance levels matching those in [12].

### Analysis of naturally occurring PL^pro^ mutants and PL^pro^ homologues with PF-07957472

Concomitant to our discovery efforts, we used publicly available sequencing information from the Global Initiative on Sharing All Influenza Data (GISAID) to understand the mutational profile of the PL^pro^ domain to assess its viability for drug discovery. PL^pro^ is one of eight structured domains on Nsp3 and is not cleaved as an independent protein (**Figure 4A and B**). For this reason, we examined the mutational frequency of all residues on Nsp3, encompassing 1944 residues, with a focus on those with mutation frequency ≥1%. This work identified 8 mutations in Nsp3 (**Figure 4B**) and the two most frequent mutations were found in the Ubl1 and SUD domains (∼66% mutation rate for both T24I and G489S). Taking a closer look at the PL^pro^ domain specific mutations, only 3 naturally occurring mutations were identified with a frequency >0.5%: P985S (0.52%), T1004I (1.02%), and V1069I (3.94%) (**Figure 4C**), all of which are outside the inhibitor binding pocket (frequencies in **Table S8**). This provides good confidence that this is a stable binding pocket and that **4** would be a stable and effective molecule against SARS-CoV-2 infections. This finding is not surprising as native substrates of PL^pro^ such as ISG15, overlap with the inhibitor binding pocket (**Figure 4D**).

**Figure 4.**
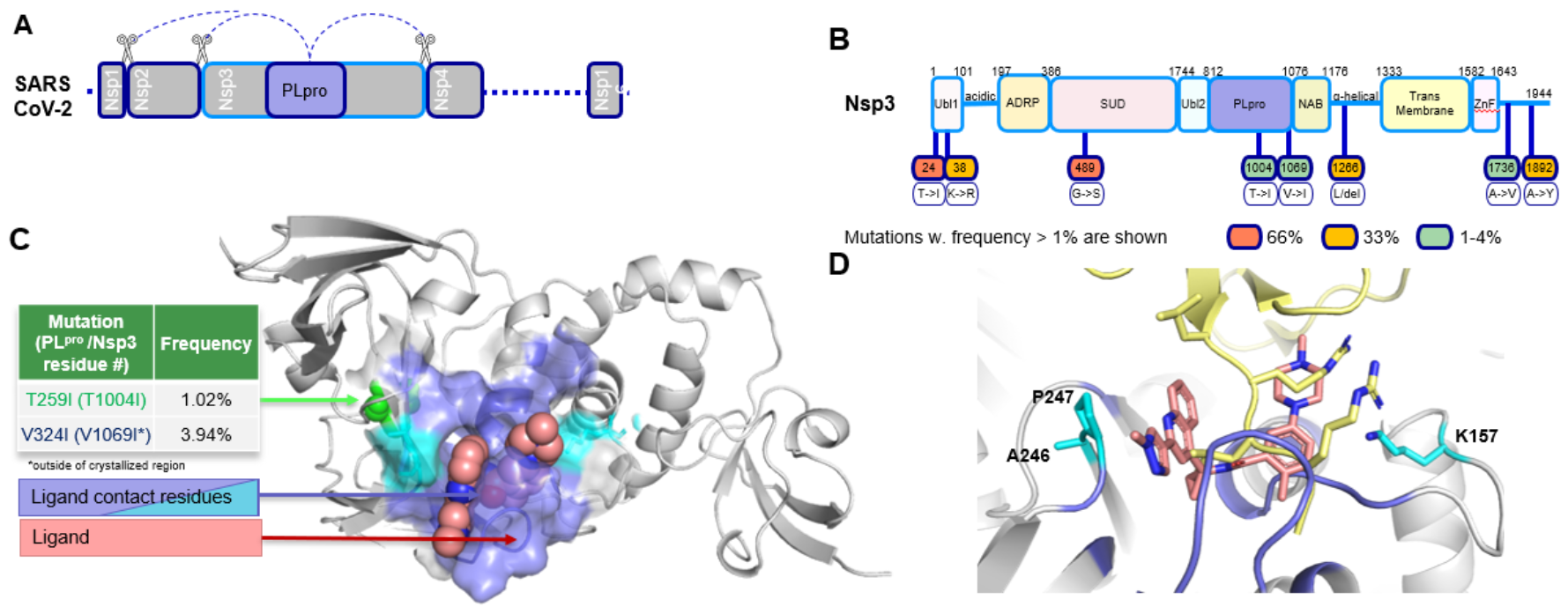
High frequency mutations in Omicron variants (6,891,950 reads) on Nsp3 and resistance selection mutants of PL^pro^. (A) The location of the cysteine protease, PL^pro^ in SARS-CoV-2 genome, and the cleavage sites it processes to generate functional viral proteins. (B) The Nsp3 protein (residues 1-1944) showing all domains identifying clinically observed Omicron mutations with frequency > 1%. (C) PL^pro^ domain (residues using PL^pro^ numbering 3-315) with location of mutation with frequency >0.5% in green which are distal from residues within 5.5 Å for **4** (pink spheres). Binding surface is shown in as blue and cyan, where cyan indicates the three highest frequency mutations (A246, P347 and K157) within the ligand binding site (**Table S8**) (D) Structural superposition of ISG15 binding site and **4** showing interaction with similar residues.

Activities of **4** were profiled against several PL^pro^ proteins from other viruses. Compound **4** was equally potent on SARS-CoV-1, but nearly inactive on other viruses like MERS, 229E and OC43 (**Table S3**), suggesting that differences in the binding pocket and especially residues in the BL2 loop (**Figure S3**) that make key interactions with the inhibitor could prevent ligand binding and cause loss of inhibitor fidelity.

## Discussion

We disclose the discovery of potent, selective, and orally available PL^pro^ inhibitors which show potent cellular antiviral activity and *in vivo* efficacy. Our results demonstrate that PL^pro^ is an essential and druggable antiviral target. Further, the pharmacokinetics and *in vitro* safety profile of our lead compound PF-07957472 (**4**) show that it is a valuable in vitro and in vivo tool for further study of PL^pro^-targeted therapeutics. More broadly, our work highlights the utility of machine learning-enabled medicinal chemistry. Through judicious use of parallel medicinal chemistry libraries, we were able to progress the campaign steadily, leading to the discovery of a compound with robust *in vivo* efficacy in less than 8 months of medicinal chemistry efforts.

## Material and Methods

See Supporting Information for detailed description of the in vitro pharmacology studies, preclinical pharmacology studies, in vivo pharmacology, off-target pharmacology, X-ray crystallography, and synthetic methods.

“All procedures performed on animals were in accordance with regulations and established guidelines and were reviewed and approved by an Institutional Animal Care and Use Committee or through an ethical review process.”

## Acknowledgments

The authors would like to acknowledge the help of Natasha Catlin and Jean Gerard Sathish.

## Data and materials availability

All data are available in the main text or the supplementary materials.

