## Supporting Information for "Discovery of SARS-CoV-2 papain-like protease (PL^pro^) inhibitors with efficacy in a murine infection model"

[1] Pfizer Research and Development
[2] PostEra PostEra, 1 Broadway, 14th floor, MA 02142, USA
[3] WuXi, WuXi AppTec (Shanghai) Co., Ltd. Shanghai, 200131
[4] Institute for Antiviral Research, Department of Animal, Dairy, and Veterinary Sciences, Utah State University; Logan, UT 84322, USA.

### In Vitro Pharmacology

#### PLpro Enzymatic Assay

The compounds were tested in a papain-like protease (PLpro) biochemical assay to determine their ability to inhibit the enzyme's activity. Biochemical potencies are reported as the concentration of the compound required to achieve 50% inhibition of the enzyme activity in the assay ($IC_{50}$). Ki values were fit to the tight binding Morrison equation with fixed parameters for enzyme concentration, substrate concentration and the Km parameter.
The potency of compounds against the SARS-CoV-2 papain-like protease (PLpro) was measured using a synthetic profluorogenic substrate, Z-RLRGG-AMC (GenScript). Compounds were serially diluted over 11 points using 100% DMSO as a diluent, from either a top concentration of 3 mM with a half-log dilution factor or a top dose of 0.1 mM with a 2-fold dilution factor. The final top dose of compounds in the assay was either 30 µM or 1 µM (1% DMSO). The assay buffer contained 50 mM HEPES, pH 7.5, 1 mM TCEP and 0.1% BSA. Final assay conditions included 6.25 nM PLPro enzyme and 25 µM peptide substrate in 1% DMSO, with an enzyme reaction time of 60 minutes.
Briefly, 250 nL of serially diluted compound was spotted into a white 384-well plate, followed by addition of SARS-CoV-2 PLpro enzyme (7.8 nM, 20 µL) in assay buffer. The compounds and PLpro enzyme were preincubated for 30 minutes at room temperature. The reaction was then initiated by the addition of profluorogenic peptide substrate in assay buffer (5 µL, 125 µM Z-RLRGG-AMC). The reaction was allowed to progress for 60 minutes at 25 °C after

which the plate was read on a Molecular Devices Spectramax M2e reader at an Ex/Em of 360 nm/460 nm. The no compound, zero percent inhibition (ZPE) control wells contained 1% DMSO with substrate and PLpro enzyme. The hundred percent effect (HPE) wells contained an internal Pfizer control compound at a dose sufficient to accomplish complete inhibition (1% DMSO), substrate and PLpro enzyme. Data were analyzed with ActivityBase software (IDBS, Ltd). The raw data were transformed to percent activity values using the average from the ZPE and HPE control wells. The resulting data were fit with the four-parameter logistic fit model to determine the $IC_{50}$ value. For compounds eliciting high potencies, the percent activity values can be fit to the Morrison equation to derive $K_i$ values with the following fixed parameters: enzyme concentration, 6.25 nM; substrate Km, 962 μM; substrate concentration, 25 μM. To qualify inter-experimental performance, the internal control (R)-5-(aminomethyl)-2-methyl-N-(1-(naphthalen-1-yl)ethyl)benzamide (compound 2[1]) was tested in each run.

##### Cellular Antiviral Activity

The ability of compounds to inhibit viral induced cytopathic effect (CPE) against human coronaviruses was measured using the VeroE6 assay with CPE and cytotoxicity endpoints using the procedure described in Dafydd R. Owen et al.[2]

Select compounds were also evaluated using SARS-CoV-2 strain USA-WA1/2020 (BEI Resources, Cat# NR-52281) in differentiated normal human bronchial epithelial (dNHBE) cells (MatTek Corporation, Ashland, MA). The cells were grown in MatTek's proprietary culture medium (AIR-100-MM) according to manufacturer's protocol and infected in a BSL-3 facility as described by Owen, et al.[2] Briefly, both virus and compound were diluted in AIR-100-MM media. Cells were infected at an approximate MOI of 0.001 for 2 hours. Inhibitor was serially diluted in media from 1 μM to 0.06 μM. Compound and virus in media was applied to the apical side whereas compound and media only was added to the basal side. Following 2 hours of infection, the apical medium was removed and washed twice with PBS and fresh compound in media was added to the basal side. At 3 days post infection, mucus was collected from the apical side and virus was quantified by plaque assay adapted from Natekar et al.[3] Briefly, infected Vero E6 cells (ATCC Cat# CRL-1586) were overlayed with semisolid overlay media (0.6% Tragacanth-Gum (Sigma, Cat# G1128-100G) and 2X-DMEM (Gibco, Cat# 11935046)), and detected using crystal violet (1% w/v) in 20% ethanol post 72 hr incubation. To determine the EC50 and EC90, the pfu/ml values were normalized to that of no drug control as a percentage of inhibition and plotted against compound concentration in GraphPad Prism software by using four-parameter logistic regression.

##### Bacterial Reverse Mutation Assay

The mutagenic potential of PF-07957472 was evaluated using the protocol described in Dafydd R. Owen et al.[2] PF-07957472 was not mutagenic or clastogenic in all tested *in vitro* genetic toxicity studies.

#### Methodology for Multi-Parameter Optimization Scoring

Tracking over time the product between cellular antiviral activity ($EC_{50}$) and metabolic CLint,app estimated in human hepatocyte incubations, was inspired by eq. 3 in J. Med. Chem. 2020, 63, 12, 6423–6435.[4]

$$\text{dose} = \frac{C_{\text{avg,ss,unbound}} \cdot CL_{\text{int}} \cdot \text{tau}}{f_a} \quad (3)$$

where the average steady-state unbound concentration ($C_{avg,ss,unbound}$) is presumed to be predictive of efficacy, $CL_{int}$ represents the intrinsic clearance, tau is the dose interval, and $f_a$ is the fraction absorbed. For the compounds in the series leading to PF-07957472, the simplifying assumption $f_a = 1$ was made, since compounds were generally permeable, and well absorbed in preclinical rodent experiments. With this assumption the estimated score is directly proportional to ($C_{avg,ss,unbound}$ * $CL_{int}$). Additionally, the desired $C_{avg,ss,unbound}$ was set to the in vitro measure of antiviral potency measure ($EC_{50}$). Nonspecific binding contributions to $CL_{int}$ and $EC_{50}$ were assumed to be minimal.

It is important to note that eq. 3 assumes a different PK-PD relationship to the trough-based efficacy relationship assumed for this project, requiring maintaining the unbound $C_{min}$ above the antiviral $EC_{90}$. However, we view eq. 3 as a generally useful score applicable to compound design if careful consideration is made to limitations of the $C_{avg,ss,unbound}$ based approach on a given project.

#### Setting Up the Collaboration Workflow and Operations

Operationally, we designed the campaign to be resilient during a pandemic: to ensure rapid design-iteration and data feedback, we set up a robust logistical workflow using the global network of CROs, and existing Pfizer internal infrastructure. Biochemical and Tier 1/2 ADME assays were replicated externally and co-located with synthesis at an external CRO, while compounds were shipped immediately post-synthesis to a BSL3 level facility for antiviral assays. This setup also ensured compounds could efficiently enter gating viral assays post-registration, and quickly progress to further characterization. The exigencies of a pandemic showcase that the decentralized drug discovery model can be an effective way to achieve operational resilience and business continuity.

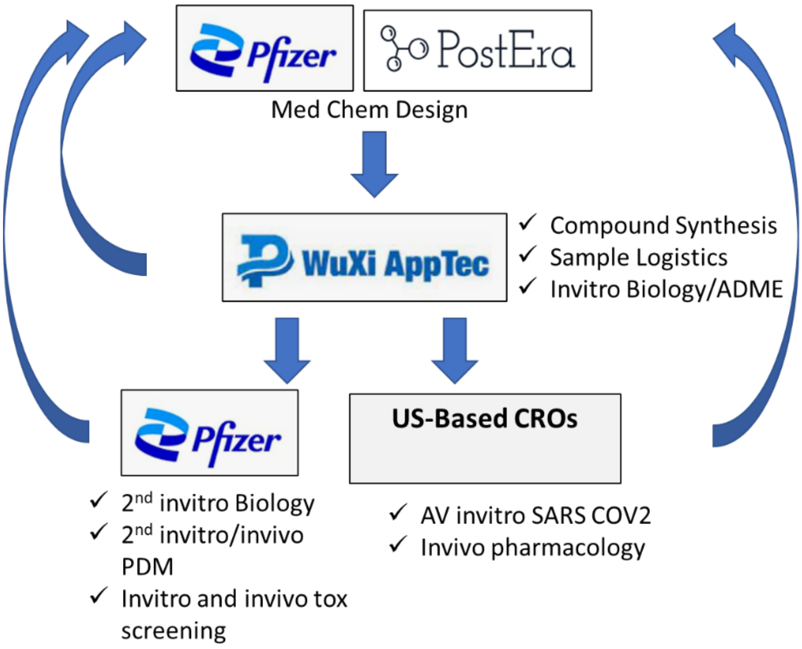

**Figure S1.** Setting Up the Collaboration Workflow and Operations

#### Preclinical Pharmacokinetics Studies

All activities involving animals were carried out in accordance with federal, state, local and institutional guidelines governing the use of laboratory animals in research in AAALAC accredited facilities and were reviewed and approved by Pfizer's or Bioduro's Institutional Animal Care and Use committee.

##### Mouse Pharmacokinetics

Mouse pharmacokinetic studies were done at BioDuro Pharmaceutical Product Development Inc. (Shanghai, PRC); male C57BL6 mice were purchased from Vital River (Beijing, China) and were typically 7-9 weeks of age at the time of dosing. During the pharmacokinetic studies all animals were housed individually. Access to food and water was provided ad libitum (i.e., subjects were dosed in the fed state). Compounds were administered intravenously (iv) via tail vein (n=2) dosed as a solution (1 mg/kg, 3 mL/kg) or via oral gavage as a solution (10 mg/kg, 3 mL/kg and 30 mg/kg, 3mL/kg) or as a suspension (15 mg/kg, 10 mL/kg; 50 mg/kg, 10 mL/kg; and 150 mg/kg, 10mL/kg). Dosing solutions were prepared immediately before dosing. The composition of each dosing vehicle is provided in Tables S5 and S6. Isoflurane was administered as anesthesia at 1-2% dose prior to collecting blood samples. Serial blood samples were collected via the retroorbital capillary at predetermined timepoints after dosing. Animals were monitored for pain or distress throughout the study, with at least daily monitoring during normal husbandry prior to study start. At the completion of the study, animals were euthanized by overdose of inhaled carbon dioxide. Blood samples were collected into tubes containing $K_2$EDTA and stored on ice until centrifugation to obtain plasma, which was stored frozen at -20 °C or lower until bioanalysis.

#### Rat Pharmacokinetics

Rat pharmacokinetic studies were done at BioDuro Pharmaceutical Product Development Inc. (Shanghai, PRC); jugular vein-cannulated male Wistar-Hannover rats were purchased from Vital River (Beijing, China) and were typically 7-9 weeks of age at the time of dosing. During the pharmacokinetic studies all animals were housed individually. Access to food and water was provided ad libitum (i.e., subjects were dosed in the fed state). In the instances where oral (po) dose was administered in the fed state, with ad libitum access to food and water. Compounds were administered intravenously (iv) via the tail vein (n = 2) dosed as a solution (1 mg/kg, 3 ml/kg) or via oral gavage as a solution (10 mg/kg, 10 ml/kg). Dosing solutions were prepared immediately before dosing. The composition of each dosing vehicle is provided in Table S5. Serial blood samples were collected via the jugular vein cannula at predetermined timepoints after dosing. Animals were monitored for pain or distress throughout the study, with at least daily monitoring during normal husbandry prior to study start. At the completion of the study, animals were euthanized by overdose of inhaled carbon dioxide. Blood samples were collected into tubes containing K2EDTA and stored on ice until centrifugation to obtain plasma, which was stored frozen at -20 °C or lower.

#### Dog Pharmacokinetics

Dog pharmacokinetic studies were done at Pfizer (Groton, CT). All procedures performed on beagle dogs were in accordance with regulations and established guidelines and were reviewed and approved by an Institutional Animal Care and Use Committee through an ethical review process. Male Beagle dogs were purchased from Marshall Bioresources (North Rose, NY); subjects 3 years of age were used in pharmacokinetics studies. For each study (n=2), compounds were dosed iv by cephalic vein as a solution (0.5 mg/kg, 1 mL/kg) or via oral gavage as a suspension (3 mg/kg, 5 mL/kg). In the instances where oral (po) dose was administered in the fed state, dogs were fed a couple hours before dosing. In the instances where the IV dose was administered in the fasted state, dogs were fasted over night and fed 4 hours post dose. Subjects were monitored for pain or distress throughout the study followed by at least daily monitoring while off study. The iv dosing vehicle was optimized such that the compounds were in solution and stable for at least 24 h. Serial blood samples were collected via the jugular vein at predefined time points post-dose. Blood samples were collected into $K_3EDTA$ treated collection tubes and were stored on wet ice prior to being centrifuged to obtain plasma, which was stored frozen at -20 °C or lower.

#### Monkey Pharmacokinetics

Monkey pharmacokinetic studies were conducted at Pfizer (Groton, CT). All procedures performed on Cynomolgus monkeys were in accordance with regulations and established guidelines and were reviewed and approved by an Institutional Animal Care and Use Committee through an ethical review process. Male Cynomolgus monkeys were purchased from Envigo Global Services (Indianapolis, IN); subjects 5-8 years of age were used in pharmacokinetic studies. For each study (n=2), compounds were dosed iv by the cephalic vein (typically 1 mg/kg, 2 ml/kg). Subjects were monitored for pain or distress throughout the study followed by at least daily monitoring while off study. The iv dosing vehicle was optimized such that the compounds were in solution and stable for at least 24 h. Serial blood samples were collected via the femoral vein at predefined time points post-dose. Blood samples were collected into $K_3EDTA$ treated collection tubes and were stored on wet ice prior to being centrifuged to obtain plasma, which was stored frozen at -20 °C or lower.

#### LC-MS/MS Analysis of Plasma Samples

Plasma samples were processed using protein precipitation with 50:50 acetonitrile: methanol containing terfenadine (5 ng/ml) as an internal standard followed by quantitation against a standard curve (0.1-1000 ng/ml) prepared in blank plasma. Quantitation of analyte in plasma samples was done using LC-MS/MS. Standard and quality control samples, prepared in blank plasma were extracted in the same manner as the in-life samples. Briefly, a Waters ACQUITY ultra performance liquid chromatography system (Waters, Milford, MA) coupled to an AB Sciex 5500 mass spectrometer equipped with an electrospray ionization source was used. Chromatographic separation was accomplished using a Kinetex C18 100A (2.6 µm, 3.00 × 50 mm) column maintained at room temperature. The mobile phase (2 solvent gradient) was optimized to achieve good separation between the analytes. Typically, solvent A consisted of 0.1% formic acid and 5 mM ammonium acetate in water, and solvent B included 0.1% formic acid and acetonitrile. The gradient generally began at 15% B until about 1.8 min, followed by an increase to 95% B to 2.21 min, then decreased to 15% B until 3 min. Analyst 1.6.1 software was used for peak integration and standard curve regression.

#### Pharmacokinetic Analysis

Pharmacokinetic parameters were calculated using noncompartmental analysis (Watson v.7.5, Thermo Scientific). The area under the plasma concentration-time curve from $t$ = 0 to infinity ($AUC_{0-\infty}$) was estimated using the linear trapezoidal rule.
Plasma clearance ($CL_p$) was calculated as:

$$CL_p = \frac{Dose_{iv}}{AUC_{0-\infty}}$$

The terminal rate constant ($k_{el}$) was calculated by linear regression of the terminal phase of the log-linear concentration-time curve and the terminal elimination $t_{1/2}$ was calculated as:

$$t_{\frac{1}{2}} = \frac{0.693}{k_{el}}$$

Apparent steady state distribution volume ($Vd_{ss}$) was determined by clearance multiplied by mean residence time. Oral bioavailability (F) was defined as:

$$F = \frac{AUC_{po} \times Dose_{iv}}{AUC_{iv} \times Dose_{po}}$$

#### Plasma Protein Binding Determination for PF-07957472

On the day of each incubation, fresh blood in K2EDTA was collected from male Wistar-Hannover rats (n=5, pooled), male Cynomolgus monkeys (n=2, pooled), and humans (n=1 male and n=1 female, pooled). The blood was centrifuged for 10 min at 2500 x g and the plasma fraction was harvested. Plasma was spiked with a final concentration of 0.3, 1, 3, or 10 µM of PF-07957472 (**4**) (final organic 1% DMSO). Fraction unbound in plasma ($f_{u,p}$) was determined by equilibrium dialysis using an HTD 96 device assembled with 12-14k molecular weight cutoff membranes (HTDialysis, LLC, Gales Ferry, CT). Dialysis chambers were loaded with 150 µl plasma and 150 µl PBS in the donor and receiver chambers, respectively. The dialysis plate was sealed with a gas-permeable membrane and stored in a

37 °C water-jacketed incubator maintained at 75% relative humidity and 5% $CO_2$, on a 100 rpm plate shaker. After a 6-hour incubation, samples were matrix-matched and quenched by protein precipitation, followed by LC-MS/MS analysis (see LC-MS/MS Analysis of Plasma Samples section above). A set of satellite samples was included to measure stability after a 6-hour incubation. Incubations were conducted with 12 replicates per concentration. $F_{u,p}$ was calculated by dividing the analyte concentration in the buffer sample by the signal in the donor sample, corrected for any dilution factors. All incubations had >70% analyte recovery and >70% stability in 6 h.

##### CYP Inhibition Assay for PF-07957472

The effects of PF-07957472 on the CYP-mediated metabolism of selective probe substrates were characterized using human liver microsomes (pool of 50 male and female donors, purchased from XenoTech (Kansas City, Kansas)).[5] Incubations (200 *µ*l) were conducted in 100 mM of potassium phosphate buffer (pH 7.4) containing 3.3 mM of $MgCl_2$, 1.2 mM of NADPH, 0.03 mg/ml microsomal protein, and probe substrates at 37 °C in a dry heat bath. A series of inhibitor stock solutions were prepared at 100-times the final test concentration. Microsomes were prewarmed with inhibitors for 5 minutes prior to initiating the reactions by the sequential addition of NADPH followed immediately by substrate. A cocktail of probe substrates was prepared at 10-times the final incubation concentration, which approximated the $K_M$: phenacetin (30 *µ*M, CYP1A2), amodiaquine (1.66 *µ*M, CYP2C8), diclofenac (6.45 *µ*M, CYP2C9), mephenytoin (39.3 *µ*M, CYP2C19), dextromethorphan (1.81 *µ*M, CYP2D6), and midazolam (2.09 *µ*M, CYP3A4). The final solvent concentration in the incubation was <1%. Reactions were terminated after 6 minutes by quenching 175-*µ*l of incubation mixture into 200 *µ*l of acetonitrile containing a cassette of internal standards (a stable labeled internal standard that was specific to each analyte was used). Following vortex mixing and centrifugation at 2300 x *g* for 5 minutes, the resulting supernatants (350 *µ*l) were transferred to clean 96-well plates, evaporated under a stream of warm nitrogen, and reconstituted in 100 *µ*l of 90/10 water/acetonitrile followed by LC-MS/MS analysis. LC-MS/MS methods and instrumentation have been previously described.

#### In Vivo Pharmacology

##### Mouse-Adapted SARS-CoV-2 Infection and Treatment Studies

The in vivo infection studies were performed in an animal biosafety level 3 (ABSL3) facility in the AAALAC-accredited Laboratory Animal Research Center at Utah State University. Pharmacokinetics studies were performed in an animal biosafety level 2 (ABSL2) facility. The study procedures were conducted with approval by the Institutional Animal Care and Use Committee at Utah State University. A total of 48 BALB/c mice (Charles River, 8 week old female) were divided into six groups as following: untreated, uninfected (mock) (n = 6), untreated infected (vehicle) (n = 6); 20 mg/kg PF-07957472 (n = 6); 50 mg/kg PF-07957472 (n = 6), 150 mg/kg PF-07957472 (n = 12). 1000 mg/kg Nirmatrelvir (n = 12). Another satellite group of six mice consisting of 2 mice/dose of PF-07957472 was treated with indicated doses (20, 50, and 150 mg/kg PF-07957472, 1000 mg/kg Nirmatrelvir, and plasma were collected at 1, 3, 6, and 12 hr post treatment for pharmacokinetic analysis. For infections, mice were anesthetized by intraperitoneal (i.p.) injection of ketamine/xylazine (50 mg/kg/5 mg/kg) and inoculated intranasally (i.n.) with 1 x $10^5$ 50% cell culture infectious dose

($CCID_{50}$) of SARS-CoV-2 MA10 (90 ml/nares). The mouse adapted MA10 virus was provided by Professor Ralph Baric (University of North Carolina).[6] For oral (p.o.) administration, PF-07957472 was solubilized in 2% (v/v) Tween80 in 98% (v/v) of 0.5% (w/v) methylcellulose in deionized water by geometric dilution. Mice were dosed twice daily (BID) x 4 days beginning at 4 hours post infection. Mice were weighed daily starting at day 0 until end of study to measure infection-associated weight loss. At 4 days post infection (dpi), mice were euthanized by isoflurane inhalation. The lungs were collected and placed in 1 ml PBS and stored at –80 °C for evaluation of lung virus titers or collected for histopathology as described below. For virus titer assays, serial $log_{10}$ dilutions of 1.0 ml lung tissue homogenates were performed in quadruplicate on confluent monolayers of Vero E6 cells seeded in 96-well microplates. The cells were incubated at 37 °C and 5% $CO_2$ for 6 days and then scored for cytopathic effect (CPE) using a light microscope. Virus lung titer ($CCID_{50}$/ml ($Log_{10}$) was calculated by linear regression using the Reed-Muench method.[7]

#### Lung Immunohistochemistry Assessment

For immunohistochemistry staining of SARS-CoV-2 nucleocapsid protein, 4 mm sections were obtained from formalin-fixed, paraffin-embedded lung tissue and immunostained using the Leica Biosystems Bond automated stainer (performed at Histowiz, Inc., Brooklyn). Epitope retrieval was performed using citrate-based pH 6 solution for 20 min at 95 ºC for heat-induced epitope retrieval (HIER). The tissue sections were then incubated with background eraser blocking reagent (Leica Biosystems) for 10 minutes to prevent non-specific binding. Next, the tissue sections were incubated for 30 min with SARS-CoV-2 (COVID-19) nucleocapsid antibody [HL448] (GTX635686, GeneTex) at a 1:10,000 dilution, followed by incubation for 30 minutes with DAB rabbit secondary reagents (Bond Polymer Refine Detection, DS9800, Leica Biosystems). The slides were visualized using an Aperio AT2 slide scanner (Leica Biosystems).

#### Lung Histopathology Assessment

To assess virus-induced damage to the lungs of SARS-CoV-2 MA10-infected mice, mice were euthanized at 4 dpi and lung lobes were collected for virus titer evaluation or left lobes were fixed in 4% paraformaldehyde at 4 °C for histopathology.[2] Fixed lung lobes were shipped to an external histology laboratory (Histowiz, Inc) for processing and blinded evaluation by an experienced veterinary pathologist and yielded similar results. Group samples (n = 6) were processed as one H&E-stained slide from each lung specimen. Each lung sample was evaluated using a semi-quantitative analysis using four parameters: perivascular inflammation, bronchial or bronchiolar epithelial degeneration or necrosis, bronchial or bronchiolar inflammation, and alveolar inflammation. A 5-point scoring system for assessment of epithelial degeneration/necrosis and inflammation was utilized (0-with normal limits; 1-mild; scattered cell necrosis/vacuolation, few/scattered inflammatory cells, 2-moderate; multifocal vacuolation or sloughed/necrotic cells, thin layer of inflammatory cells, 3-marked; multifocal/segmental necrosis, epithelial loss/effacement, thick layer of inflammatory cells, 4-severe; coalescing areas of necrosis; parenchymal effacement, confluent areas of inflammation. A total pathology score was calculated for each mouse by adding the individual histopathological scores.

##### Statistics and Figures

All graphs were generated using GraphPad Prism. The statistical analysis was performed as follows. For the body weight, the % initial data from day 1 to 4, each dose group was compared to placebo group (0 mg/kg PF-09757472) via mixed model analysis. Raw (unadjusted) p-values are reported. For the lung virus $log_{10}$ titer, and histopathology score data each dose group was compared to that of the 0 mg/kg group via one-way ANOVA. Raw (unadjusted) p-values are reported.

#### Off Target Pharmacology

##### Mammalian Protease Panel

These standard assays were run at Reaction Biology Corporation, except for cathepsin F at BPS Bioscience.

##### Coronavirus Protease Panel

A set of compounds was tested against a Coronovirus protease panel (SARS-CoV-1, SARS-CoV-2, 229E, MERS, and OC43), as described in the "PLPro Enzymatic Assay" section above, to determine their ability to inhibit each enzyme's activity.[8-13] The potency of compounds against each protease was measured using a synthetic profluorogenic substrate, Z-RLRGG-AMC (GenScript). Compounds were serially diluted over 11 points using 100% DMSO as a diluent, from a top concentration of 1 mM with a half-log dilution factor. The final top dose of compounds in the assay was 10 μM (1% DMSO). The assay buffer contained 50 mM HEPES, pH 7.5, 1 mM TCEP and 0.1% BSA. Final assay conditions were enzyme-specific and are noted below:

| **Assay** | **Enzyme (nM)** | **Substrate (μM)** | **$K_m$ (μM)** |
|---|---|---|---|
| hCoV-229E_PLpro2 | 100 | 31.25 | 253.7 |
| MERS_Plpro | 100 | 25 | 639.5 |
| hCoV-OC43_PLpro2 | 100 | 31.25 | 1671 |
| SARS-CoV-1_Plpro | 6.25 | 25 | 282 |
| SARS-CoV-2_Plpro | 6.25 | 25 | 962 |

Briefly, 250 nL of serially diluted compound was spotted into a white 384-well plate, followed by addition of 20 μL of enzyme in assay buffer. The compounds and PLpro enzyme were preincubated for 30 minutes at room temperature. The reaction was then initiated by the addition of profluorogenic peptide substrate in assay buffer (5 μL, Z-RLRGG-AMC). The reaction was allowed to progress for 60 minutes (SARS-CoV-1, SARS-CoV-2, and OC43), 80 minutes (MERS), or 90 minutes (229E) at 25 °C, after which the plate was read on a Molecular Devices Spectramax M2e reader at an Ex/Em of 360 nm/460 nm.

##### Data Analysis for Mammalian and Coronavirus Protease Panels

Data were analyzed with ActivityBase software (IDBS). The no compound, zero percent inhibition (ZPE) control wells contained 1% DMSO with substrate and enzyme. The hundred percent effect (HPE) wells contained a control compound at a dose sufficient to accomplish complete inhibition (1% DMSO), substrate and enzyme.  The raw data were transformed to percent activity values using the average from the ZPE and HPE control wells. The resulting data were fit with the four-parameter logistic fit model to determine the $IC_{50}$ value. Ki values were fit to the tight binding Morrison equation with fixed parameters for enzyme concentration, substrate concentration and the Km parameter using ActivityBase software (IDBS).

$$\mathrm{TB\,Ki}: v_i = b + v0*\left(1 - \frac{2*[I]}{E+[I]+Ki*\frac{[S]+Km}{Km}+\sqrt{(E+[I]+Ki*\frac{[S]+Km}{Km})^2-4E[I]}}\right) = \mathrm{function}(b, v0, Ki, E)$$

#### Ion Channels

**Compound Preparation**

Compounds were dissolved and initially diluted in dimethyl sulfoxide (DMSO), with a final dilution in external solution to generate final working concentrations. The final DMSO concentration in all experiments was 0.33% (v/v). All tissue culture media and reagents were obtained from Thermo Fisher (Waltham, MA, USA), unless otherwise stated.
Ion Channel experiments were performed at Metrion Biosciences Ltd (Cambridge, UK).

**hERG**

Chinese hamster ovary (CHO) cells stably expressing the human $K_v$11.1 (hERG) channel (hERG DUO, B'SYS GmbH, Witterswil, Switzerland) were cultured in Ham's F12 + Glutamax medium supplemented with 10% (v/v) fetal bovine serum (FBS), 0.1 mg/mL Geneticin, 0.1 mg/mL Hygromycin B and 10 mM HEPES. Cells were cultured at 37°C and 5% $CO_2$ and subcultured every 2-3 days in T175 flasks. On the day of the experiment, cells were harvested at ~80% confluency by rinsing with HBSS ($Ca^{++}$/$Mg^{++}$ -/-) and incubating in 2 mL TrypLE (Gibco) for 3-4 min at 37°C. To quench TrpLE, 8 mL of Hams F-12+ 10% FBS was added and cells were triturated to create a single cell suspension. Cells were centrifuged at 1000 rcf for 2 min and resuspended in CHO-S-SFM II (Gibco) at a density of 3 million cells per mL.

Ionic currents were evaluated in the whole-cell configuration using the Qube384 automated planar patch clamp platform (Sophion Bioscience A/S, Ballerup, Denmark). QChip 384X plates, containing 10 patch clamp holes per well, were used to maximize success rate, which was routinely > 95%. For hERG experiments, the external solution was composed of (in mM): 140 NaCl, 2 KCl, 2 $CaCl_2$, 1 $MgCl_2$, 10 HEPES, 5 Glucose, pH 7.4, 350 mOsM. The internal solution contained (in mM): 20 KCl, 10 EGTA, 10 HEPES, 120 KF, pH 7.2, 335 mOsM. The hERG current was elicited from a holding potential of -80 mV by a voltage step to +40 mV for 500 ms, followed by a repolarizing ramp to -80 mV at -0.6 mV/ms. This pattern was repeated at a rate of 0.05 Hz. Peak hERG current was measured during the ramp. All studies were conducted at 23 °C.

Three vehicle periods each lasting 5 minutes were applied to establish a stable baseline. This was followed by the addition of increasing concentrations of test compound, with each exposure lasting 5 minutes. Patch clamp data were analyzed using Analyzer Software 6.4.72 (Sophion Bioscience A/S, Ballerup, Denmark). Current amplitudes were determined by averaging the last 4 currents under each test condition. The percentage inhibition of each compound was determined by taking the ratio of current amplitude measured in the presence of various concentrations of the test compound ($I_{Compound}$) versus the vehicle control current ($I_{Vehicle}$): % Inhibition = [1-($I_{Compound}$/$I_{Vehicle}$)] * 100%. A dose-response curve was generated and fit to the Hill equation by the Sophion Analyzer software to determine an $IC_{50}$

value for each compound. The minimum and the slope of the fit were free fitted, with the top being fixed to 100% inhibition.

**hCav1.2**
Human embryonic kidney (HEK) cells stably expressing human Cav1.2/β2a/α2δ1 calcium channel (SB Drug Discovery Glasgow, UK, hCav1.2 clone #10) were cultured in DMEM medium supplemented with 10% FBS, Geneticin (0.6 mg/ml), hygromycin (0.1 mg/ml), and blasticidin (0.004 mg/ml). The calcium channel antagonist verapamil hydrochloride (0.025 mg/ml) was added to culture media to minimize calcium-induced cytotoxicity. Cells were cultured at 37°C and 5% $CO_2$ and subcultured every 3-4 days in T175 flasks. 24 hours prior to cell harvesting, culture media was supplemented with 1mM Na-butyrate to enhance channel expression. On the day of the experiment, cells were harvested at ~80% confluency by rinsing with DPBS ($Ca^{++}$/$Mg^{++}$ -/-) and incubating in 2 mL TrypLE (Gibco) for 3-4 min at 37°C. To quench TrpLE, 8 mL of Hams F-12+ 10% FBS was added and cells were triturated to create a single cell suspension. Cells were centrifuged at 1000 rcf for 2 min and resuspended in CHO-S-SFM II (Gibco) at a density of 3 million cells per mL.

Ionic currents were evaluated in the whole-cell configuration using the Qube384 automated planar patch clamp platform using QChip 384X plates. For hCav1.2 experiments, the external solution was composed of (in mM): 137.9 NaCl, 5.3 KCl, 10 HEPES, 0.49 $MgCl_2$, 10 $CaCl_2$, 5.5 glucose, 4.16 $NaHCO_3$, 0.34 $Na_2HPO_4$, 0.41 $MgSO_4$, pH 7.4, 315 mOsM. The internal solution contained (in mM): 90 CsCl, 50 CsF, 2 $MgCl_2$, 10 EGTA, 10 HEPES, 2 NaCl, 2 Na-ATP, pH 7.2, 307 mOsM. The hCav1.2 current was elicited from a holding potential of -40 mV by a voltage step to 0 mV for 150 ms. This pattern was repeated at 0.05 Hz, and hCav1.2 current was measured as the peak current at 0 mV. All studies were conducted at 23 °C.

Three vehicle periods each lasting 5 minutes were applied to establish a stable baseline. This was followed by the addition of increasing concentrations of test compound, with each exposure lasting 5 minutes. Patch clamp data were analyzed using Analyzer Software 6.4.72. Current amplitudes were determined by averaging the last 4 currents under each test condition. The percentage inhibition of each compound was determined by taking the ratio of current amplitude measured in the presence of various concentrations of the test compound ($I_{Compound}$) versus the vehicle control current ($I_{Vehicle}$): % Inhibition = $[1-(I_{Compound}/I_{Vehicle})] * 100\%$. A dose-response curve was generated and fit to the Hill equation by the Sophion Analyzer software to determine an $IC_{50}$ value for each compound. The minimum and the slope of the fit were free fitted, with the top being fixed to 100% inhibition.

**hNav1.5**
CHO cells stably expressing the human Nav1.5 sodium channel (University of Pennsylvania, Philadelphia, PA, hNav1.5 clone #6) were cultured in Hams F12 + Glutamax media supplemented with 10% FBS and Geneticin (0.5 mg/mL) at 37°C and 5% $CO_2$ and subcultured every 2-3 days in T175 flasks. On the day of the experiment, cells were harvested at ~80% confluency by rinsing with HBSS ($Ca^{++}$/$Mg^{++}$ -/-) and incubating in 2 mL TrypLE (Gibco) for 3-4 min at 37°C. To quench TrpLE, 8 mL of Hams F-12+ 10% FBS was added and cells were triturated to create a single cell suspension. Cells were centrifuged at 1000 rcf for 2 min and resuspended in CHO-S-SFM II (Gibco) at a density of 3 million cells per mL.

Ionic currents were evaluated in the whole-cell configuration using the Qube384 platform using QChip 384x plates. For hNav1.5 experiments, the external solution was composed of (in mM): 140 NaCl, 5 KCl, 3 $CaCl_2$, 1.2 $MgCl_2$, 5 HEPES, 11.1 glucose, pH 7.4, 305 mOsM. The internal solution contained (in mM): 120 CsF, 20 CsCl, 15 NaCl, 10 HEPES, 10 EGTA, pH 7.2, 285 mOsM. For the hNav1.5 current, from an initial holding potential of -80 mV, a 200 ms prepusle to -120 mV was used to homogenize channel inactivation, followed by a 40 ms step to a test potential of -15 mV. Membrane potential was futher depolarized to +40 mV for 200 ms, followed by a ramp from +40 mV to -80 mV (-1.2 mV/ms). This voltage pattern was repeated at 0.2 Hz, wit the hNav1.5 peak current defined as the maximum current during the step to -15 mV. All studies were conducted at 23 °C

Three vehicle periods each lasting 5 minutes were applied to establish a stable baseline. This was followed by the addition of increasing concentrations of test compound, with each exposure lasting 5 minutes. Patch clamp data were analyzed using Analyzer Software 6.4.72. Current amplitudes were determined by averaging the last 4 currents under each test condition. The percentage inhibition of each compound was determined by taking the ratio of current amplitude measured in the presence of various concentrations of the test compound ($I_{Compound}$) versus the vehicle control current ($I_{Vehicle}$):   % Inhibition = $[1-(I_{Compound}/I_{Vehicle})] * 100\%$.  A dose-response curve was generated and fit to the Hill equation by the Sophion Analyzer software to determine an $IC_{50}$ value for each compound. The minimum and the slope of the fit were free fitted, with the top being fixed to 100% inhibition.

#### CEREP in vitro pharmacology panel PF-07957472 Screening

Secondary (off-target) pharmacology studies were conducted by Eurofins Cerep (Celle-Lévescault, France) on behalf of Pfizer, Inc. The in vitro off-target pharmacology of PF-07957472 was assessed at 30 μM in a broad target profiling panel which represents targets with known links to potential safety concerns and includes G-protein coupled receptors, ion channels, transporters, and enzymes according to established protocols.

**Table S1.  % response of PF-07957472 in CEREP in vitro off-target safety panel at 30 μM**

| Functional Agonism | % response | Functional Antagonism | % response |
|---|---|---|---|
| Adenosine A1 | -1 | Adrenergic Alpha 1a | 17 |
| Adenosine A2a | 0 | Adrenergic Alpha 2b | 11 |
| Adrenergic Alpha 1a | -1 | Adrenergic Beta 1 | -31 |
| Adrenergic Alpha 2a | -1 | Adrenergic Beta 2 | -52 |
| Adrenergic Alpha 2b | 3 | Angiotensin 1 | 12 |
| Adrenergic Beta 1 | 1 | Cannabinoid 1 | 10 |
| Adrenergic Beta 2 | 12 | Dopamine 1 | -24 |
| Angiotensin 1 | -2 | Histamine 1 | 10 |
| Cannabinoid 1 | 10 | Muscarinic 1 | 13 |
| Cholecystokinin 2 | 5 | Muscarinic 2 | 30 |
| Dopamine 1 | 1 | Muscarinic 3 | 30 |
| EndothelinA | 2 | Opioid Mu | 6 |
| Histamine 1 | -2 | CRF1 | -35 |

| | | | |
|---|---|---|---|
| Histamine 2 | 4 | MC2R | -28 |
| Histamine 3 | -10 | TRH1 | 38 |
| Muscarinic 1 | 2 | **Enzyme** | **% inhibition** |
| Muscarinic 2 | -1 | Angiotensin Converting Enzyme | 14 |
| Muscarinic 3 | -4 | Acetylcholinesterase | 17 |
| Neurokinin 1 | -3 | Cyclooxygenase 2 | 36 |
| Opioid Delta | 0 | Monoamine Oxidase | 16 |
| Opioid Kappa | -9 | PDE3B | 4 |
| Opioid Mu | 22 | PDE4D2 | 20 |
| Serotonin 1a | 0 | **Kinase** | **% inhibition** |
| Serotonin 1b | 8 | Abl Kinase | 2 |
| Serotonin 2a | 0 | Aurora A Kinase | -1 |
| Serotonin 2b | 5 | EGFR Kinase | 9 |
| Serotonin 4e | -3 | Lck Kinase | -1 |
| Vasopressin 1a | 1 | P38 MAP Kinase | 10 |
| CRF1 | 2 | Src Kinase | -5 |
| MC2R | 1 | VEGFR2 Kinase | 11 |
| TRH1 | 2 | **Transporters-Binding** | **% inhibition** |
| **Ion Channel- Binding** | **% inhibition** | Norepinephrine Transporter | -6 |
| L-Type Ca (verapamil) | -19 | Dopamine Transporter | 8 |
| L-Type Ca (Nifedipine) | -6 | Serotonin Transporter | 8 |
| L-Type Ca (Diltiazem) | 10 | Choline Transporter | 25 |
| | | GABA Transporter | -4 |
| GABAa1 Receptor | 10 | **NHR-Binding** | **% inhibition** |
| GABAa (benzodiazepine Site) | 7 | Androgen Receptor (Binding) | -10 |
| AMPA Receptor | 13 | Glucocorticoid Receptor (Binding) | 2 |
| NMDA Receptor | -2 | PPAR gamma (Binding) | 9 |
| NMDA Receptor (PCP site) | 14 | | |
| Nicotinic Ach Receptor (muscle) | 2 | | |
| Nicotinic Ach Receptor (Neuronal) | -6 | | |
| Serotonin 3 | -10 | | |
| Sodium (Site 2) | -15 | | |

Abbreviations: GPCR, G-Protein Coupled Receptor; NHR, Nuclear Hormone Receptor; EGFR, Endothelial Growth Factor Receptor; VEGFR, Vascular Endothelial Growth Factor Kinase; Lck, Lymphocyte-Specific Protein Tyrosine Kinase; MAPK14, Mitogen Activated Protein Kinase 14 (p38α).

### X-ray Crystallography

#### SARS-CoV-2 PL$^{pro}$ protein purification and co-crystallization with PF-07957472

The codon-optimized gene corresponding to SARS-CoV-2 PL$^{pro}$ domain (Nsp3 residues 1562-1878; Uniprot P0DTD1; PL$^{pro}$ domain numbering 1-315) bearing a cysteine to serine mutation in the catalytic triad (C111S) was synthesized as a gBlock gene fragment by Integrated DNA Technologies and was subcloned into a pET28a vector using NcoI and XhoI restriction enzymes (New England Biolabs). The protein was expressed with a cleavable N-terminal $His_{6x}$ tag in BL21(DE3) *E. coli* cells (Novagen Singles$^{TM}$). The protein was isolated from the lysate using Ni-NTA Agarose resin (Qiagen), the $His_{6x}$ tag was cleaved using tobacco etch virus protease (TEV), and the tag-free construct was purified to homogeneity and >95% purity by size exclusion chromatography using a HiLoad Superdex 75 pg 16/600 column (Cytiva) in a buffer containing 50 mM HEPES pH 8, 200 mM NaCl, and 1 mM TCEP. After concentrating the protein to 10 mg/mL, PF-07957472 was added to the protein at a final concentration of 300 µM. Crystals of the protein-ligand complex were obtained in a crystallization condition containing 0.1 M Sodium citrate tribasic dihydrate pH 5.0, 30% v/v Jeffamine® ED-2001 pH 7.0.

#### Diffraction Data, Structure Determination and Refinement

| | |
|---|---|
| Resolution range | 68.51 - 2.595 (2.688 - 2.595) |
| Space group | P 21 21 21 |
| Unit cell | 64.866 99.662 188.682 90 90 90 |
| Total reflections | 496277 (52163) |
| Unique reflections | 37135 (229) |
| Multiplicity | 13.4 (13.7) |
| Completeness (%) | 79.10 (5.28) |
| Mean I/sigma(I) | 7.75 (0.53) |
| Wilson B-factor | 64.80 |
| R-merge | 0.1912 (2.701) |
| R-meas | 0.199 (2.831) |
| R-pim | 0.05448 (0.8239) |
| CC1/2 | 0.993 (0.976) |
| CC* | 0.998 (0.994) |

| | |
|---|---|
| Reflections used in refinement | 30588 (201) |
| eflections used for R-free | 1480 (7) |
| R-work | 0.2371 (0.4539) |
| R-free | 0.2808 (0.5321) |
| CC(work) | 0.872 (0.433) |
| CC(free) | 0.867 (0.199) |
| Number of non-hydrogen atoms | 6770 |
| macromolecules | 6532 |
| ligands | 140 |
| solvent | 164 |
| Average B-factor | 66.94 |
| macromolecules | 67.35 |
| Ligands | 54.71 |
| Solvent | 56.22 |

X-ray crystal diffraction data were collected on the IMCA 17-ID beamlines (Advanced Photo Source, Argonne National Laboratory) using a wavelength of 0.95374 Å. All datasets were processed with autoPROC (Global Phasing).[14] Anisotropy was corrected by ellipsoidal truncation using STARANISO as a part of autoPROC (Global Phasing) [ref: Tickle, I.J., *et al.* STARANISO. Cambridge, UnitedKingdom: Global Phasing Ltd. 2018.]. Phasing for each structure was obtained by molecular replacement in PHASER[15] using as search model 7CJM. The crystals obtained of SARS-CoV-2 PL$^{pro}$ in complex with PF-07957472 were in the P212121 space group with three protomers in the asymmetric unit. Inhibitor density was strongest in two PL$^{pro}$ protomers, likely owing to crystal packing at the compound binding site. Manual model building was performed in COOT[16] and refinement using BUSTER (Global Phasing) [ref: Bricogne G., *et al.* BUSTER version X.Y.Z. Cambridge, United Kingdom: Global Phasing Ltd. 2017.] Molecular graphics were generated using PyMOL. Data collection and refinement statistics are provided in Table 1.

**Table S2.** Data collection and refinement statistics.

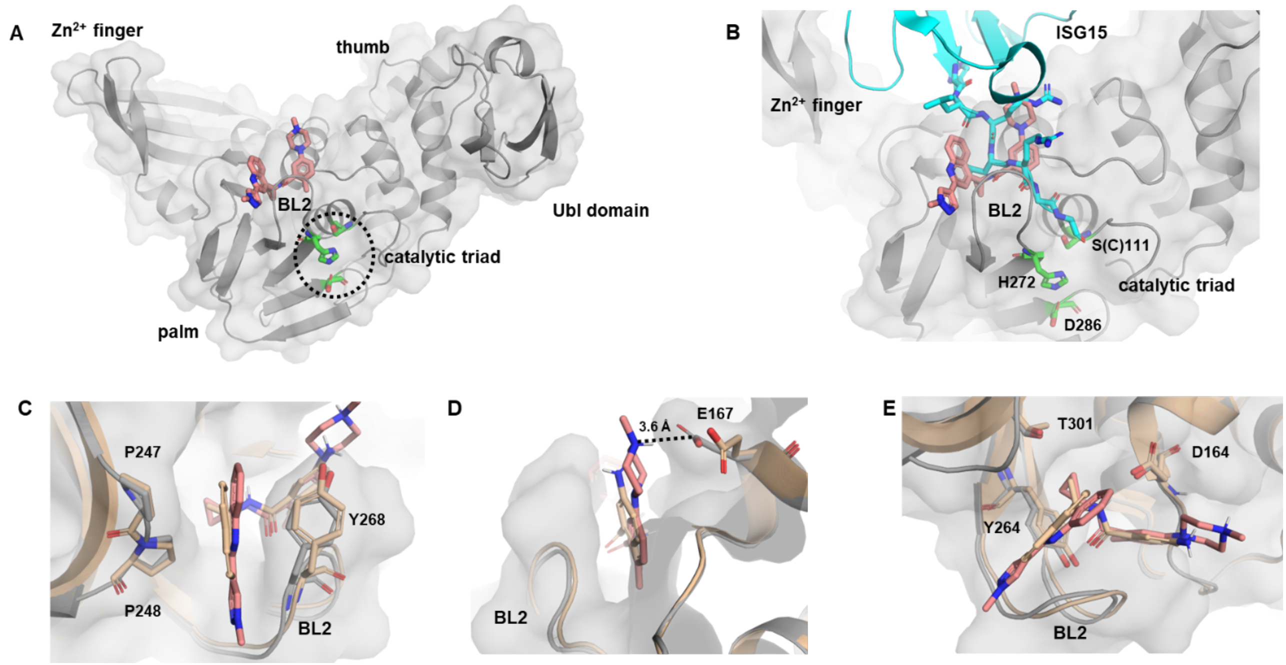

**Figure S2**. Interactions formed between SARS-CoV-2 PL$^{pro}$ and PF-07957472, and comparisons with the GRL0617-SARS-CoV-2 PL$^{pro}$ complex. (**A**) The X-ray crystal structure of SARS-CoV-2 PL$^{pro}$ (grey cartoon and surface representation) bound to PF-07957472 (pink) with subdomains and the BL2 loop labelled, and the catalytic triad shown in green. (**B**)

Structural superimposition of SARS-CoV-2 $PL^{pro}$ bound to PF-07957472 (grey/pink) and SARS-CoV-2 $PL^{pro}$ bound to mouse ISG15 (cyan; PDB ID 6YVA). The $PL^{pro}$ molecule from 6YVA is not shown for clarity. Seven residues at the C-terminus of mISG15, which overlap with the compound binding site, are shown in stick representation (His149 – Gly155). The catalytic triad is shown in green; a catalytically dead construct of PLpro (C111S) was used to generate the crystal structure. (**C-E**) Structural overlays of SARS-CoV-2 PLpro bound to PF-07957472 (grey/pink) and SARS-CoV-2 bound to GRL0617 (sand-colored; PDB ID 7CMD). The orientation of the binding pocket varies in each panel, with the BL2 loop labelled throughout. Surface representation is shown for the structure of SARS-CoV-2 $PL^{pro}$ bound to PF-07957472 only (grey).

### Materials and Methods: Synthetic Procedures

#### Abbreviations

| | |
|---|---|
| M | Molar |
| MeOD-$d_4$ | Deuterated methanol |
| $N_2$ | Nitrogen gas |
| $Na_2SO_4$ | Sodium sulfate |
| $NaBH_3CN$ | Sodium cyanoborohydride |
| $NaHCO_3$ | Sodium bicarbonate |
| NaOH | Sodium hydroxide |
| Pd(dppf)Cl2 | [1,1′-Bis(diphenylphosphino)ferrocene]dichloropalladium (II) |
| $Pd(PPh_3)_2Cl_2$ | Palladium(II)bis(triphenylphosphine) dichloride |
| prep-HPLC | Preparative High Performance Liquid Chromatography |
| sat. | saturated |
| aq. | aqueous |
| TEA | Triethylamine |
| DMF | Dimethylformamide |
| min | minute |
| h | hour |
| m-CPBA | meta-chloroperoxybenzoic acid |
| LCMS | Liquid Chromatography Mass Spectrometry |
| HCl | Hydrochloric acid |

| | |
|---|---|
| HATU | N-[(Dimethylamino)-1H-1,2,3-triazolo-[4,5-b]pyridin-1-ylmethylene]-N-methylmethanaminium hexafluorophosphate N-oxide |
| $POCl_3$ | Phosphorus(V) oxychloride |
| $Na_2CO_3$ | Sodium carbonate |
| CbzOSu | N-(benzyloxycarbonyloxy)succinimide |

#### General Synthetic Methods

All chemicals, reagents, and solvents were purchased from commercial sources and were used without further purification. 1H NMR data are reported relative to residual solvent signals and are reported as follows: chemical shift (ppm), multiplicity, coupling constant (Hz), and integration. The multiplicities are denoted as: s, singlet; d, doublet; t, triplet; q, quartet; m, multiplet; br s, broad singlet. Silica gel chromatography was performed using Biotage or ISCO purification systems with pre-packaged columns. Concentration under reduced pressure was performed on a rotary evaporator with a water bath temperature not exceeding 60 ℃. Purity of final compounds was assessed by HPLC with UV detection at 215 or 254 nm; all tested compounds showed > 90% purity.

#### Synthesis and Characterization of Compounds 2-4

##### Synthesis of Compound 2

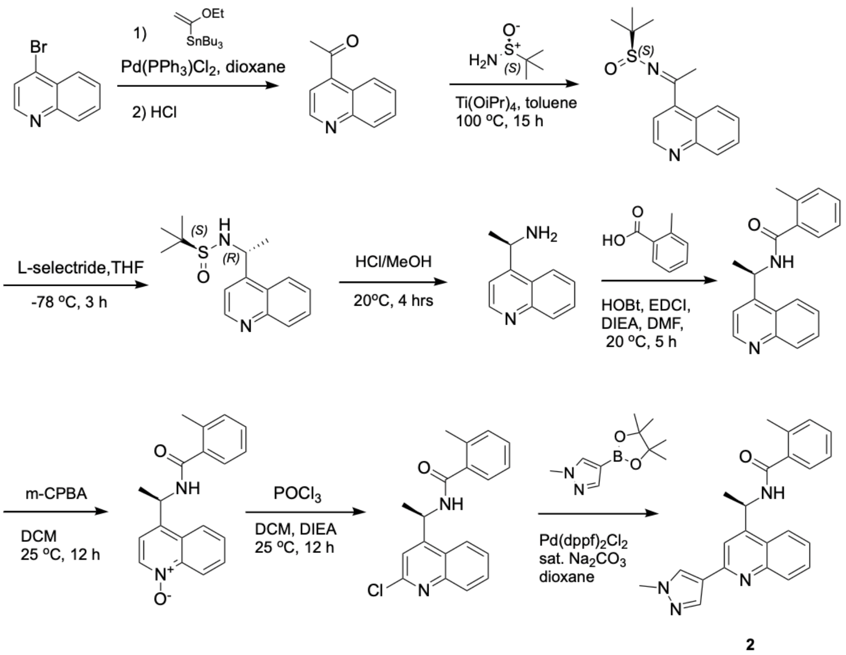

2

**(*S*)-2-methyl-N-((*R*)-1-(quinolin-4-yl)ethyl)propane-2-sulfinamide**.

**Step 1.** To a solution of 4-bromoquinoline (20000.0 mg, 96.128 mmol) in 1,4-dioxane (500.0 mL) under $N_2$ were added tributyl(1-ethoxyvinyl)stannane (68030.0 mg, 188.37 mmol) and $Pd(PPh_3)_2Cl_2$ (6750 mg, 9.61 mmol) at 20 ˚C. The reaction mixture was stirred at 100 ˚C under $N_2$ for 15 h. LCMS showed a peak with a desired mass. The reaction mixture was quenched with 200 mL sat. aq. KF at 0 ˚C then partitioned between ethyl acetate and water

(500/500 mL). The organic layer was separated and the aqueous layer was re-extracted with ethyl acetate (500 ml). The combined organic layer was washed with brine, dried and evaporated *in vacuo* to give a residue. The residue was dissolved in 1,4-dioxane (500 mL) containing conc. HCl (50 mL) at 0 °C. The mixture was stirred at 20 °C for 15 h before being evaporated in vacuo to afford a crude. The crude was dissolved in 200 mL methanol and basified with ammonium hydroxide to pH = 9 at 0 °C. The mixture was then evaporated in vacuo and purified by silica gel chromatography using a Biotage (330 g column, petroleum ether/ethyl acetate = 1/0 to 0/1) to afford 1-(quinolin-4-yl)ethan-1-one (14850 mg, 90%) as a yellow oil.

**Step 2.** To a solution of 1-(quinolin-4-yl)ethan-1-one (20300.0 mg, 118.58 mmol) in toluene (500.0 mL) under $N_2$ were added (*S*)-2-methylpropane-2-sulfinamide (43100 mg, 356 mmol) and Ti(O-iPr)$_4$ (135000 mg, 474 mmol ) at 20 °C. The reaction mixture was stirred at 100 °C under $N_2$ for 15 h. LCMS showed a peak with a desired mass. The reaction mixture was then evaporated in vacuo to afford a crude, which was purified by silica gel chromatography using a Biotage (330 g column, petroleum ether/ethyl acetate = 1/0 to 0/1) to afford (*S*)-2-methyl-N-(1-(quinolin-4-yl)ethylidene)propane-2-sulfinamide (32500 mg) as a yellow oil.

**Step 3.** To a solution of (*S*)-2-methyl-N-(1-(quinolin-4-yl)ethylidene)propane-2-sulfinamide (32000 mg, 116.63 mmol) in THF (500 mL) under $N_2$, was added L-selectride (210.0 mL, 1 M, 210 mmol) dropwise at -78 °C. The reaction mixture was stirred at -78 °C under $N_2$ for 5 h before being quenched by 100 mL sat. aq. $NH_4Cl$ at 0 °C. The reaction mixture was then partitioned between ethyl acetate and water (500/500 mL). The organic layer was separated and the aqueous layer was re-extracted with ethyl acetate (500 mL). The combined organic layer was washed with brine, dried and evaporated *in vacuo* before being purified by silica gel chromatography using a Biotage (330 g column, petroleum ether/ethyl acetate = 1/0 to 0/1, ethyl acetate/methanol = 1/0 to 0/1) to afford (*S*)-2-methyl-N-((*R*)-1-(quinolin-4-yl)ethyl)propane-2-sulfinamide (24560 mg, 76%) as a colorless gum. LCMS *m/z* calc. for $C_{15}H_{21}N_2OS^+$ $[M+H]^+$ 277.1, found 277.0.

**(*R*)-1-(quinolin-4-yl)ethan-1-amine**. To a solution of (*S*)-2-methyl-N-((*R*)-1-(quinolin-4-yl)ethyl)propane-2-sulfinamide (14.5 g, 52 mmol) in MeOH (100 mL) was added HCl/MeOH (100 mL, 400 mmol, 4M in MeOH). The reaction was stirred at 20 °C for 4 h. TLC (petroleum ether/ethyl acetate = 3/1) showed that starting material was consumed and a new spot was detected. The mixture was filtered and the filter cake was dissolved in water (20 mL). The solution was basified with sat. aq. $NaHCO_3$ to pH = 8 and extracted with dichloromethane/MeOH (v/v = 10/1, 5 × 200 mL). The organic layers were combined, dried over $Na_2SO_4$, filtered, evaporated under reduced pressure to give (*R*)-1-(quinolin-4-yl)ethan-1-amine (6 g, 66%) as a yellow oil. $^1$H NMR (400 MHz, MeOD-d$_4$) δ 1.52 (d, *J* = 6.63 Hz, 3H), 4.93 (q, *J* = 6.71 Hz, 1H), 7.53-7.69 (m, 2H), 7.73-7.85 (m, 1H), 8.05 (d, *J* = 8.50 Hz, 1H), 8.20 (d, *J* = 8.51 Hz, 1H), 8.82 (d, *J* = 4.63 Hz, 1H). LCMS *m/z* calc. for $C_{11}H_{13}N_2^+$ $[M+H]^+$ 173.1, found 173.1.

**(*R*)-2-methyl-*N*-(1-(quinolin-4-yl)ethyl)benzamide**. To a solution of (*R*)-1-(quinolin-4-yl)ethan-1-amine (2.8 g, 16.26 mmol) in DMF (28 mL) were added 2-methylbenzoic acid (2.66 g, 19.51 mmol), HOBt (7.4 g, 19.51 mmol) and DIPEA (4.2 g, 32.52 mmol) at 20 °C. The mixture was cooled to 0 °C before adding EDCI (3.74 g, 19.51 mmol). The reaction was then stirred at 20 °C for 5 h. TLC (PE/EA = 1/1.5) showed that starting material was consumed and new spots were detected. Water (200 mL) was then added to the reaction, and the mixture was extracted with ethyl acetate (6 × 100 mL). The organic layers were combined, washed with brine, filtered and concentrated under reduced pressure to give a residue, which was purified by column chromatography on a silica gel column (petroleum ether/ethyl acetate = 100/0 to 1/1) to give (*R*)-2-methyl-*N*-(1-(quinolin-4-yl)ethyl)benzamide (3 g, 64%) as white solid. $^1$H NMR (400 MHz, $CDCl_3$) δ 1.73 (d, *J* = 6.75 Hz, 3H), 2.41 (s, 3H), 6.01-6.12 (m, 1H), 6.27 (br d, *J* = 7.88 Hz, 1H), 7.11-7.24 (m, 2H), 7.27 (s, 2H), 7.40 (d, *J* = 4.50 Hz, 1H), 7.62 (ddd, *J* = 8.32, 7.00, 1.19 Hz, 1H), 7.74 (td, *J* = 7.63, 1.25 Hz, 1H), 8.13 (d, *J* = 8.50 Hz, 1H), 8.22 (d, *J* = 8.38 Hz, 1H), 8.85 (d, *J* = 4.38 Hz, 1H). LCMS *m/z* calc. for $C_{19}H_{19}N_2O^+$ $[M+H]^+$ 291.1, found 291.1.

**(*R*)-4-(1-(2-methylbenzamido)ethyl)quinoline 1-oxide.** To a solution of (*R*)-2-methyl-*N*-(1-(quinolin-4-yl)ethyl)benzamide (2.5 g, 8.61 mmol) in dichloromethane (25 mL) was added m-CPBA (2.62 g, 12.92 mmol) portion-wise with stirring at 0 °C. The reaction was warmed to 25 °C and stirred at 25 °C for 12 h. TLC (petroleum ether/tetrahydrofuran = 1/1) showed that starting material was consumed and a new spot was detected. The reaction was quenched with sat. aq. $NaHCO_3$ (100 mL). The organic layer was washed with brine, dried over $Na_2SO_4$, filtered and concentrated under reduced pressure to give a residue, which was purified by column chromatography on a silica gel column (eluting with petroleum ether/tetrahydrofuran = 100/1 to 0/1) to give (*R*)-4-(1-(2-methylbenzamido)ethyl) quinoline 1-oxide (2.5 g, 95%) as a white solid. $^1$H NMR (400 MHz, $CDCl_3$) δ ppm 1.77 (br d, *J* = 6.75 Hz, 3H), 2.47 (s, 3H), 5.88-6.14 (m, 1H), 7.09 (br d, *J* = 6.13 Hz, 1H), 7.17-7.25 (m, 2H), 7.28-7.34 (m, 2H), 7.39 (br d, *J* = 7.38 Hz, 1H), 7.38-7.62 (m, 2H), 8.18 (br d, *J* = 8.25 Hz, 1H), 8.28 (br d, *J* = 8.88 Hz, 2H). LCMS *m/z* calc. for $C_{19}H_{19}N_2O_2^+$ $[M+H]^+$ 307.1, found 307.2.

**(*R*)-*N*-(1-(2-chloroquinolin-4-yl)ethyl)-2-methylbenzamide.** To a solution of (*R*)-4-(1-(2-methylbenzamido)ethyl)quinoline 1-oxide (1.5 g, 4.9 mmol) and DIPEA (6.33 g, 48.96 mmol) in dichloromethane (50 mL) was added $POCl_3$ (2.8 g, 24.48 mmol) dropwise with stirring at 0 °C. The reaction was warmed to 25 °C and stirred at 25 °C for 12 h. TLC (THF) showed that starting material was consumed and new spots were detected. The reaction was transferred to a cooled sat. aq. $NaHCO_3$ solution (100 mL) and extracted with dichloromethane (3 × 50 mL). The combined organic layer was washed with brine, dried over $Na_2SO_4$, filtered and concentrated under reduced pressure to give a residue, which was purified by column chromatography on a silica gel column (eluting with petroleum ether/tetrahydrofuran = 100/1 to 3/1) to give (*R*)-*N*-(1-(2-chloroquinolin-4-yl)ethyl)-2-methylbenzamide (0.6 g , 38%) as a white solid. $^1$H NMR (400 MHz, $CDCl_3$) δ 1.70-1.85 (m, 3H), 2.45 (s, 3H), 5.95-6.17 (m, 2H), 7.18-7.26 (m, 2H), 7.31-7.37 (m, 2H), 7.42 (s, 1H), 7.60-7.70 (m, 1H), 7.74-7.84 (m, 1H), 8.07 (d, *J* = 8.00 Hz, 1H), 8.20 (d, *J* = 8.50 Hz, 1H). LCMS *m/z* calc. for $C_{19}H_{18}ClN_2O^+$ $[M+H]^+$ 325.1, found 325.1.

**(*R*)-2-methyl-*N*-(1-(2-(1-methyl-1*H*-pyrazol-4-yl)quinolin-4-yl)ethyl) benzamide (2).** To a solution of (*R*)-*N*-(1-(2-chloroquinolin-4-yl)ethyl)-2-methylbenzamide (50.0 mg, 0.154 mmol) in dioxane (4.0 mL) were added 1-methyl-4-(4,4,5,5-tetramethyl-1,3,2-dioxaborolan-2-yl)-1*H*-pyrazole (96.1 mg, 0.462 mmol), Pd(dppf)$Cl_2$ (11.3 mg, 0.0154 mmol), and sat. aq. $Na_2CO_3$ (0.5 mL) at 20 °C. The mixture was bubbled with $N_2$ for 1 min then stirred at 100 °C for another 15 h. LCMS showed a peak with desired mass. The reaction mixture was filtered and the filtrate was purified by prep-HPLC to afford (*R*)-2-methyl-*N*-(1-(2-(1-methyl-1*H*-

pyrazol-4-yl)quinolin-4-yl)ethyl) benzamide (18.8 mg, 33%) as a white solid after lyophilization.

Prep-HPLC condition:

Column: C18-1 150*30mm*5um

Mobile phase: water (formic acid)-acetonitrile; acetonitrile: 0% - 37%

Flow rate (mL/min)：30 mL/min.

1H NMR (400 MHz, DMSO-$d_6$) δ = 8.95 (d, *J* = 7.5 Hz, 1H), 8.41 (s, 1H), 8.23 (d, *J* = 8.0 Hz, 1H), 8.09 (s, 1H), 7.97 (d, *J* = 8.5 Hz, 1H), 7.86 (s, 1H), 7.73 (t, *J* = 7.5 Hz, 1H), 7.57 (t, *J* = 7.3 Hz, 1H), 7.38-7.30 (m, 2H), 7.28-7.21 (m, 2H), 5.89 (m, 1H), 3.94 (s, 3H), 2.28 (s, 3H), 1.61 (d, *J* = 7.0 Hz, 3H). LCMS *m/z* calc. for $C_{23}H_{23}N_4O^+$ $[M+H]^+$ 371.2, found 371.3.

**Synthesis of 2-methyl-5-(4-methylpiperazin-1-yl)benzoic acid (int-1):**

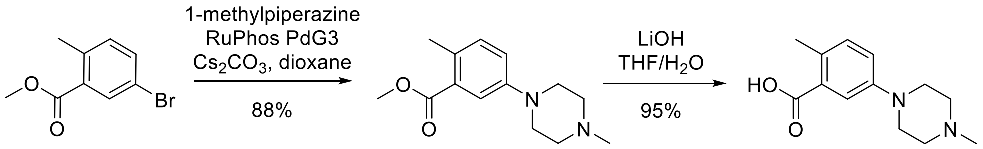

**int-1**

**Step 1.** A 500-mL 3-neck RBF equipped with overhead stirrer, internal temperature probe, and reflux condenser under nitrogen was charged with methyl 5-bromo-2-methylbenzoate (18 g, 78.6 mmol), cesium carbonate (76.8 g, 236 mmol), anhydrous dioxane (200 mL) and 1-methylpiperazine (9.6 mL, 86.4 mmol). The resulting yellow slurry was sparged with nitrogen under vigorous stirring for 15 min. The flask was briefly opened and $RuPhosPdG_3$ (3.3 g, 3.93 mmol) was added in one portion and reaction mixture sparged for another 10 min. The reaction was heated under nitrogen in an aluminium block to 100 °C for 16 hours. The cooled reaction mixture was filtered over celite, rinsing with EA. The amber filtrate (600 mL total) was concentrated to remove dioxane, then dissolved in EA (150 mL). The EA solution was washed once with half-saturated ammonium chloride (75 mL). The aqueous was extracted once with EA (100 mL). The combined organics were extracted with 1M HCl (1 x 100 mL). The dark aq. layer was made neutral by the addition of 25 mL 4M NaOH, then extracted with EA (150 mL). TLC showed desired product still present in the aq. layer, so it was made basic (pH~12) by the addition of 4 mL 4M NaOH and extracted again with EA (100 mL). By TLC, almost all of the desired product was now in the combined organics. The combined organics were washed once with saturated brine, dried over anhydrous magnesium sulfate, filtered and concentrated under reduced pressure to afford 17.98 g (92%) of methyl 2-methyl-5-(4-methylpiperazin-1-yl)benzoate as an amber oil. 1H NMR (400 MHz, $CDCl_3$) δ 7.48 (d, *J* = 2.7 Hz, 1H), 7.13 (d, *J* = 8.2 Hz, 1H), 6.99 (dd, *J* = 2.7, 8.2 Hz, 1H), 3.89 (s, 3H), 3.28 - 3.16 (m, 4H), 2.64 - 2.55 (m, 4H), 2.50 (s, 3H), 2.37 (s, 3H). LCMS *m/z* calc. for $C_{14}H_{20}N_2O_2$ [M+H]+ 249.16 found 249.6.

**Step 2.** A solution of methyl 2-methyl-5-(4-methylpiperazin-1-yl)benzoate (17.98 g, 72.4 mmol) in EtOH (54 mL) and water (18 mL) was treated with lithium hydroxide monohydrate (3.1 g, 79.6 mmol) and the reaction mixture was heated to 60 °C for 16 hrs. The reaction was cooled to rt and the amber-colored solution was treated with con HCl gradually until pH = 3 (12 mL). A solid formed and the entire reaction became a solid block of beige clay. The solid was treated with 150 mL EtOH and broken up with a spatula until a thick, but stirrable, slurry formed. The beige slurry was stirred at rt for 1.5 hrs and then filtered, rinsing the filter cake with minimal amount of ethanol. The solids were dried in high vacuo for 18 hours at rt. After overnight drying of the first crop afforded 12.8 g (65%) of 2-methyl-5-(4-methylpiperazin-1-yl)benzoic acid (**int-1**) as a white solid. 1H NMR (400 MHz, DMSO-$d_6$) δ ppm 12.82 (br s, 1H), 10.99 (br s, 1H), 7.38 (d, *J* = 2.7 Hz, 1H), 7.23 - 7.14 (m, 1H), 7.13 - 7.06 (m, 1H), 3.76 (br d, *J* = 1.6 Hz, 2H), 3.44 (br dd, *J* = 7.0, 14.0 Hz, 2H), 3.12 (br s, 4H), 2.79 (s, 3H), 2.41 (s, 3H). LCMS *m/z* calc. for $C_{13}H_{18}N_2O_2$ [M+H]+ 235.3, found 235.5

**Synthesis of Compound 3**

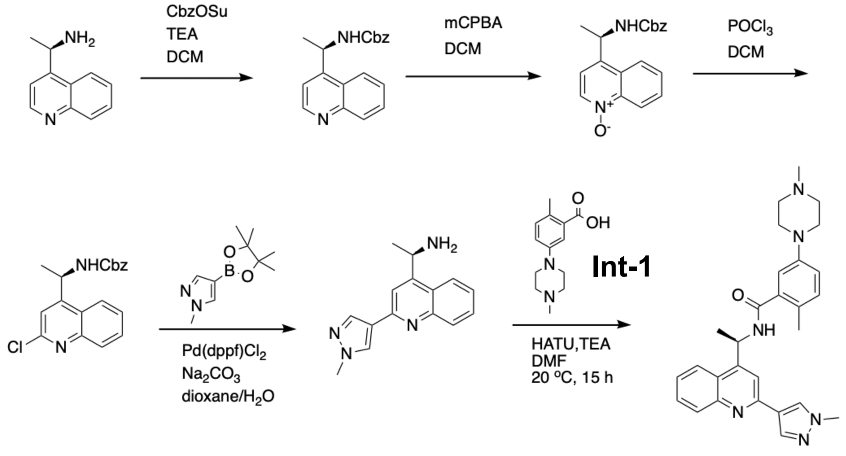

3

**Benzyl (*R*)-(1-(quinolin-4-yl)ethyl)carbamate**. To a stirred solution of (*R*)-1-(quinolin-4-yl)ethan-1-amine (8400.0 mg, 48.77 mmol) in dichloromethane (500 mL) were added TEA (7400 mg, 73.2 mmol) and CbzOSu (13400 mg, 53.7 mmol) at 0 ˚C. The reaction mixture was stirred at 25 ˚C for 2 h. LCMS showed a mass peak of the desired product. The reaction was evaporated in vacuo and purified by silica gel chromatography using a Biotage (120 g column, petroleum ether/ethyl acetate = 1/0 to 0/1) to afford benzyl (*R*)-(1-(quinolin-4-yl)ethyl)carbamate (14600 mg, 98%) as a yellow oil. LCMS *m/z* calc. for $C_{19}H_{19}N_2O_2^+$ $[M+H]^+$ 307.1, found 307.0.

**(*R*)-4-(1-(((benzyloxy)carbonyl)amino)ethyl)quinoline 1-oxide**. To a solution of benzyl (*R*)-(1-(quinolin-4-yl)ethyl)carbamate (7000.0 mg, 22.85 mmol) in dichloromethane (100 mL) was slowly added m-CPBA (6960 mg, 34.3 mmol) in several portions at 5 ˚C. After that, the ice-water bath was removed, and the mixture was warmed up to 30 ˚C and stirred for 20 h. LCMS showed a mass peak of the desired product. The reaction mixture was evaporated in vacuo and purified by silica gel chromatography using a Biotage (120 g silica column, petroleum ether/ethyl acetate = 1/0 to 0/1) to afford (*R*)-4-(1-(((benzyloxy)carbonyl)amino)ethyl)quinoline 1-oxide (5700 mg, 77%) as a yellow oil. LCMS *m/z* calc. for $C_{19}H_{19}N_2O_3^+$ $[M+H]^+$ 323.1, found 323.0.

**Benzyl (*R*)-(1-(2-chloroquinolin-4-yl)ethyl)carbamate**. To a solution of (*R*)-4-(1-(((benzyloxy)carbonyl)amino)ethyl)quinoline 1-oxide (5700.0 mg, 17.68 mmol) in dichloromethane (20.0 mL) was added $POCl_3$ (27100 mg, 177 mmol) at 0 ˚C. The reaction mixture was stirred at 25 ˚C for 16 h. LCMS showed a mass peak of the desired product. $NaHCO_3$ (80 mL sat. aq.) was then added to the reaction mixture at 25 ˚C. The aqueous phase was extracted with dichloromethane (20 mL x 2). The combined organic layer was dried with anhydrous $Na_2SO_4$, filtered, evaporated in vacuo, and purified by silica gel chromatography using a Biotage (120 g column, petroleum ether/ethyl acetate = 1/0 to 0/1) to afford benzyl (*R*)-(1-(2-chloroquinolin-4-yl)ethyl)carbamate (4960 mg, 82%) as a yellow solid. $^1$H NMR (400 MHz, DMSO-d$_6$) δ = 8.28 (d, *J* = 8.4 Hz, 1H), 8.22 (br d, *J* = 7.6 Hz, 1H), 7.99 (d, *J* = 8.3 Hz, 1H), 7.85 (t, *J* = 7.3 Hz, 1H), 7.71 (br t, *J* = 7.5 Hz, 1H), 7.50 (s, 1H), 7.43-7.19 (m, 5H), 5.76 (s, 2H), 5.56-5.45 (m, 1H), 1.47 (d, *J* = 7.0 Hz, 3H). LCMS *m/z* calc. for $C_{19}H_{18}ClN_2O_2^+$ $[M+H]^+$ 341.1, found 340.9.

**(*R*)-1-(2-(1-methyl-1*H*-pyrazol-4-yl)quinolin-4-yl)ethan-1-amine**. To a solution of benzyl (*R*)-(1-(2-chloroquinolin-4-yl)ethyl)carbamate (2000.0 mg, 5.868 mmol ) in 1,4-dioxane (50.0 mL) under $N_2$ were added 1-methyl-4-(4,4,5,5-tetramethyl-1,3,2-dioxaborolan-2-yl)-1*H*-pyrazole (2440 mg, 11.7 mmol ) and Pd(dppf)$Cl_2$ (429 mg, 0.587 mmol) and sat. aq. $Na_2CO_3$ (5.0 mL) at 20 °C. The reaction mixture was stirred at 100 °C under $N_2$ for 15 h. LCMS showed a mass peak of the desired product. The reaction mixture was evaporated *in vacuo* and purified by silica gel chromatography using a Biotage (40 g column, petroleum ether/ethyl acetate = 1/0 to 0/1, ethyl acetate/methanol = 1/0 to 0/1) to afford (*R*)-1-(2-(1-methyl-1*H*-pyrazol-4-yl)quinolin-4-yl)ethan-1-amine (606 mg, 27%) as a yellow solid. $^1$H NMR (400MHz, $CDCl_3$) δ = 8.18-8.13 (m, 1H), 8.12 (s, 1H), 8.05 (br d, *J* = 8.4 Hz, 1H), 7.97 (br d, *J* = 8.1 Hz, 1H), 7.88 (s, 1H), 7.66 (br t, *J* = 7.4 Hz, 1H), 7.47 (br t, *J* = 7.3 Hz, 1H), 5.01 (br d, *J* = 6.1 Hz, 1H), 3.94 (s, 3H), 1.65-1.51 (m, 3H). LCMS *m/z* calc. for $C_{15}H_{17}N_4^+$ $[M+H]^+$ 253.1, found 253.0. (and benzyl (*R*)-(1-(2-(1-methyl-1*H*-pyrazol-4-yl)quinolin-4-yl)ethyl)carbamate (1023 mg, 69%) as a red solid. LCMS *m/z* calc. for $C_{23}H_{23}N_4O_2^+$ $[M+H]^+$ 387.2, found 387.2).

**(*R*)-2-methyl-*N*-(1-(2-(1-methyl-1H-pyrazol-4-yl)quinolin-4-yl)ethyl)-5-(4-methylpiperazin-1-yl)benzamide (3)**. To a solution of 2-methyl-5-(4-methylpiperazin-1-yl)benzoic acid (**int-1**) (107 mg, 0.396 mmol) in DMF (10 mL) were added HATU (226 mg, 0.594 mmol), TEA (120 mg, 1.19 mmol), (*R*)-1-(2-(1-methyl-1*H*-pyrazol-4-yl)quinolin-4-yl)ethan-1-amine (100.0 mg, 0.396 mmol) at 20 °C. The reaction mixture was stirred at 20 °C for 15 h before being partitioned between ethyl acetate and water (50/50 mL). The organic layer was separated and the aqueous layer was re-extracted with ethyl acetate (50 mL). The combined organic layer was washed with brine, dried and evaporated in vacuo to afford a crude. The crude was dissolved in 5 mL DMF and purified by prep-HPLC to afford (*R*)-2-methyl-*N*-(1-(2-(1-methyl-1*H*-pyrazol-4-yl)quinolin-4-yl)ethyl)-5-(4-methylpiperazin-1-yl)benzamide (52 mg, 28%) as a yellow solid after lyophilization.
Prep-HPLC condition:
Column: Boston Prime C18 150*30mm*5um
Mobile phase: water (ammonium hydroxide) – acetonitrile:acetonitrile %: 14%-54%
Flow Rate (mL/min)：30 mL/min
$^1$H NMR (400 MHz, MeOD-d$_4$) δ ppm 8.34 (s, 1H), 8.28 (d, *J* = 7.8 Hz, 1H), 8.18 (s, 1H), 8.08 (d, *J* = 8.0 Hz, 1H), 7.88 (s, 1H), 7.78 (t, *J* = 7.5 Hz, 1H), 7.70-7.56 (m, 1H), 7.15 (d, *J* = 7.8 Hz, 1H), 7.02-6.93 (m, 2H), 6.02 (q, *J* = 6.9 Hz, 1H), 4.19 (s, 1H), 4.02 (s, 3H), 3.22-3.14 (m, 4H), 2.68-2.57 (m, 4H), 2.36 (s, 3H), 2.26 (s, 3H), 1.74 (d, *J* = 7.0 Hz, 3H). LCMS *m/z* calc. for $C_{28}H_{33}N_6O^+$ $[M+H]^+$ 469.3, found 469.5.

**Synthesis of Compound 4**

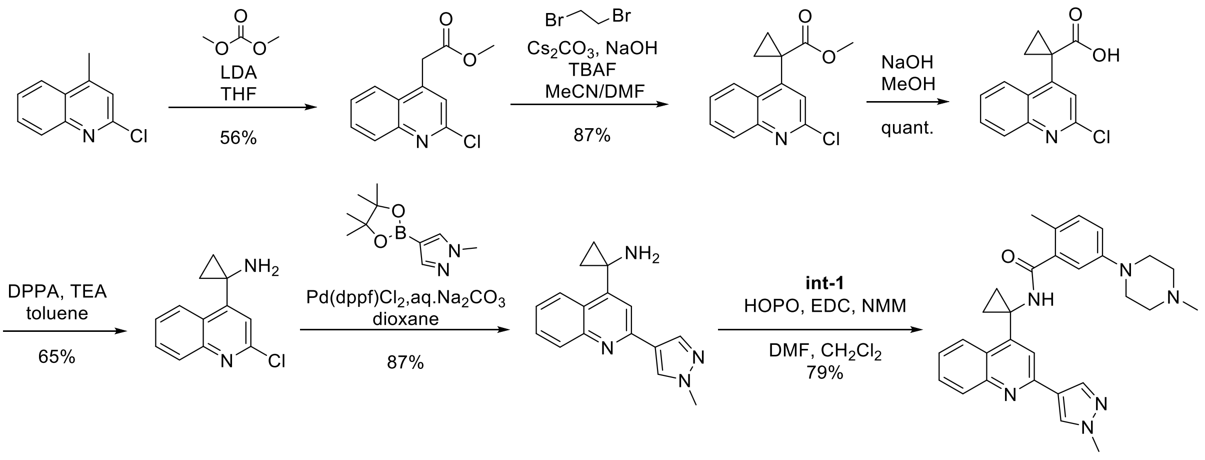

**1-(2-(1-methyl-1H-pyrazol-4-yl)quinolin-4-yl)cyclopropan-1-amine**

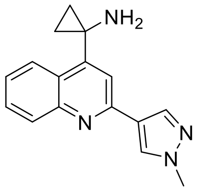

**Step 1. Preparation of methyl 2-(2-chloroquinolin-4-yl)acetate**

To a solution of 2-chloro-4-methylquinoline (1000.0 mg, 6.984 mmol) in THF (30 mL) under $N_2$ was added LDA (6.0 mL, 2 M, 12 mmol) at -78 °C. The reaction mixture was stirred at -78 °C under $N_2$ for 30 min, followed by the addition of dimethyl carbonate (1270 mg, 14.1 mmol) at 0 °C. The reaction mixture was stirred at 0 °C under $N_2$ for 1 h, then quenched with 3 mL sat. $NH_4Cl$ at 0 °C. The reaction mixture was partitioned between ethyl acetate and $H_2O$ (200/50 mL), the organic layer was separated, and the aqueous layer was re-extracted with ethyl acetate (50 mL). The combined organic layer was washed with brine, dried with anhydrous $Na_2SO_4$, and evaporated *in vacuo*. The residue was purified by silica gel chromatography using a Biotage (20 g silica gel column, PE: EA= 1: 0 to 0: 1) to afford methyl 2-(2-chloroquinolin-4-yl)acetate (744 mg, 56.1%) as a yellow oil. LCMS *m/z* calc. for $C_{12}H_{11}ClNO_2^+$ $[M+H]^+$ 236.0, found 236.1. $^1H$ NMR (400 MHz, $CDCl_3$-d) δ = 8.06 (dd, *J* = 0.8, 8.6 Hz, 1H), 7.99 - 7.93 (m, 1H), 7.76 (ddd, *J* = 1.4, 7.0, 8.4 Hz, 1H), 7.66 - 7.57 (m, 1H), 7.36 (s, 1H), 4.06 (s, 2H), 3.73 (s, 3H).

**Step 2. Preparation of methyl 1-(2-chloroquinolin-4-yl)cyclopropane-1-carboxylate**

To a solution of methyl 2-(2-chloroquinolin-4-yl)acetate (3000.0 mg, 12.73 mmol) in DMF (60.0 mL) were added 1,2-dibromoethane (3590 mg, 19.1 mmol), $Cs_2CO_3$ (8300 mg, 25.5 mmol), tetra-*n*-butylammonium fluoride (13 mL, 1 M, 13 mmol), and NaOH (1020 mg, 25.5 mmol) at 20 °C. The reaction mixture was stirred at 20 °C for 15 h. LCMS showed a mass peak of the desired product. The reaction mixture was then partitioned between ethyl acetate and $H_2O$ (250/100mL). The organic layer was separated, and the aqueous layer was re-extracted with ethyl acetate (150 mL). The combined organic layer was washed with brine, dried with anhydrous $Na_2SO_4$, filtered, and concentrated *in vacuo*. The residue was then purified by silica gel chromatography using a Biotage (80 g silica gel column, PE: EA= 1: 0 to 0: 1) to afford methyl 1-(2-chloroquinolin-4-yl)cyclopropane-1-carboxylate (2885 mg, 86.6%) as a yellow solid. LCMS *m/z* calc. for $C_{14}H_{13}ClNO_2^+$ $[M+H]^+$ 262.1, found 262.0. $^1H$ NMR (400 MHz, $CDCl_3$-d) δ = 8.10 - 7.99 (m, 2H), 7.74 (ddd, *J* = 1.4, 7.0, 8.4 Hz, 1H), 7.59 (ddd, *J*

= 1.4, 7.0, 8.4 Hz, 1H), 7.35 (s, 1H), 3.58 (s, 3H), 1.89 (br d, *J* = 3.1 Hz, 2H), 1.32 (br d, *J* = 2.3 Hz, 2H).

**Step 3. Preparation of 1-(2-chloroquinolin-4-yl)cyclopropane-1-carboxylic acid**
To a solution of methyl 1-(2-chloroquinolin-4-yl)cyclopropane-1-carboxylate (2885.0 mg, 11.02 mmol) in MeOH (60mL) was added 10% aq. NaOH (15mL) at 20 °C. The reaction mixture was stirred at 20 °C for 15 h. LCMS showed a mass peak of the desired product. The reaction mixture was then acidified with 1N HCl to pH = 1 at 0 °C and partitioned between ethyl acetate and $H_2O$ (250/50 mL). The organic layer was separated, and the aqueous layer was re-extracted with ethyl acetate (150 mL). The combined organic layer was washed with brine, dried with anhydrous $Na_2SO_4$, and evaporated in vacuo to afford 1-(2-chloroquinolin-4-yl)cyclopropane-1-carboxylic acid (2730 mg, 100%) as a yellow solid. LCMS *m/z* calc. for $C_{13}H_{11}ClNO_2^+$ $[M+H]^+$ 248.0, found 248.1. $^1H$ NMR (400 MHz, DMSO-d6) δ ppm 12.72 (br s, 1 H), 8.12 (d, J=7.81 Hz, 1 H), 7.98 (d, J=8.53 Hz, 1 H), 7.83 (ddd, J=8.39, 7.02, 1.37 Hz, 1 H), 7.68 (t, J=7.61 Hz, 1 H), 7.56 (s, 1 H), 1.66 - 1.78 (m, 2 H), 1.37 (br s, 2 H).

**Step 4. Preparation of 1-(2-chloroquinolin-4-yl)cyclopropan-1-amine**
To a solution of 1-(2-chloroquinolin-4-yl)cyclopropane-1-carboxylic acid (2500.0 mg, 10.09 mmol) in toluene (60.0 mL) under $N_2$, was added diphenylphosphoryl azide (3060 mg, 11.1 mmol) and TEA (2250 mg, 22.2 mmol) at 20 °C. The reaction mixture was stirred at 90 °C under $N_2$ for 2 h, then evaporated in vacuo to afford a crude. The crude was dissolved in 1,4-dioxane (40mL), followed by adding aq. NaOH (10%, 20 mL) at 20 °C under $N_2$. The reaction mixture was stirred at 90 °C for 15 h. LCMS showed a mass peak of the desired product. The reaction mixture was then partitioned between ethyl acetate and $H_2O$ (250/150 mL). The organic layer was separated, and the aqueous layer was re-extracted with ethyl acetate (50 mL). The combined organic layer was washed with brine, dried with anhydrous $Na_2SO_4$, filtered and concentrated *in vacuo*. The residue was dissolved in 30 mL MeOH, then 10 mL 1 N HCl was added to the solution at 20 °C. The reaction mixture was stirred at 20 °C for 1 h, then partitioned between ethyl acetate and $H_2O$ (250/150 mL). The organic layer was discarded, and the aqueous layer was basified with $NH_3.H_2O$ to pH = 9 at 0 °C, followed by extraction with DCM (200 mL). The organic layer was washed with brine, dried with anhydrous $Na_2SO_4$ and concentrated *in vacuo* to afford 1-(2-chloroquinolin-4-yl)cyclopropan-1-amine (1444 mg, 65.4%) as a white solid. LCMS *m/z* calc. for $C_{12}H_{12}ClN_2^+$ $[M+H]^+$ 219.1, found 219.1.

**Step 5. Preparation of 1-(2-(1-methyl-1*H*-pyrazol-4-yl)quinolin-4-yl)cyclopropan-1-amine**
To a solution of 1-(2-chloroquinolin-4-yl)cyclopropan-1-amine (1000.0 mg, 4.573 mmol) in 1,4-dioxane (40.0 mL) under $N_2$ were added sat. aq. $Na_2CO_3$ (8.0 mL), Pd(dppf)$Cl_2$ (335mg, 0.457 mmol), and 1-methyl-4-(4,4,5,5-tetramethyl-1,3,2-dioxaborolan-2-yl)-1*H*-pyrazole (1900 mg, 9.15 mmol) at 20 °C. The reaction mixture was stirred at 100 °C under $N_2$ for 15 h. LCMS showed a mass peak of the desired product. The reaction mixture was then filtered, and the filtrate was concentrated *in vacuo* and purified by silica gel chromatography using a Biotage (20 g silica gel column, PE: EA= 1: 0 to 0: 1) to afford 1-(2-(1-methyl-1*H*-pyrazol-4-yl)quinolin-4-yl)cyclopropan-1-amine (1055 mg, 87.3%) as a yellow oil. LCMS *m/z* calc. for $C_{16}H_{17}N_4^+$ $[M+H]^+$ 265.1, found 265.1. $^1H$ NMR (400 MHz,

METHANOL-$d_4$) δ = 8.96 (s, 1H), 8.64 (d, *J* = 8.6 Hz, 1H), 8.60 (s, 1H), 8.51 (s, 1H), 8.42 (d, *J* = 8.6 Hz, 1H), 8.17 (dt, *J* = 1.2, 7.8 Hz, 1H), 8.05 - 7.98 (m, 1H), 4.10 (s, 3H), 1.87 - 1.79 (m, 2H), 1.68 - 1.61 (m, 2H).

**PF-07957472 (4): 2-methyl-*N*-(1-(2-(1-methyl-1*H*-pyrazol-4-yl)quinolin-4-yl)cyclopropyl)-5-(4-methylpiperazin-1-yl)benzamide**

A round bottom flask under nitrogen was charged with 2-methyl-5-(4-methylpiperazin-1-yl)benzoic acid (2770 mg, 10.2 mmol), DCM (50mL), DMF (4mL), HOPO (1130 mg, 10.2 mmol) and *N*-methylmorpholine (1700 mg, 16 mmol , 1.8 mL) in that order. To this was then added 1-(2-(1-methyl-1*H*-pyrazol-4-yl)quinolin-4-yl)cyclopropan-1-amine (3000 mg, 10 mmol) followed by EDCI (2550 mg, 13.3 mmol). The reaction mixture was stirred at rt for 15 hours. The mixture was diluted with DCM (100 mL), washed with saturated $NH_4Cl$ solution, water, brine and dried over anhydrous $Na_2SO_4$. Filtration and solvent removal afforded a crude product that started to solidify (6.12 g). This was combined with 50 mL of EA and 10 mL of heptane and mixture stirred at rt for 24 hours. The white solid was collected by filtration and concentrated *in vacuo* to give 2-methyl-*N*-(1-(2-(1-methyl-1*H*-pyrazol-4-yl)quinolin-4-yl)cyclopropyl)-5-(4-methylpiperazin-1-yl)benzamide (2.69 g, 55%). $^1$H NMR (400 MHz, $CDCl_3$-d) δ ppm 8.32 (d, *J* = 8.20 Hz, 1 H), 8.21 (s, 1 H), 8.15 (s, 1 H), 8.11 (d, *J* = 8.20 Hz, 1 H), 8.01 (s, 1 H), 7.71 (t, *J* = 7.22 Hz, 1 H), 7.52 -7.57 (m, 1 H), 7.01 (d, *J* = 8.59 Hz, 1 H), 6.83 (dd, *J* = 8.39, 2.54 Hz, 1 H), 6.69 (d, *J* = 2.73 Hz, 1 H), 6.47 (s, 1 H), 4.02 (s, 3 H), 3.04-3.12 (m, 4 H), 2.48-2.57 (m, 4 H), 2.34 (s, 3 H), 2.10 (s, 3 H), 1.60-1.67 (m, 1 H), 1.43-1.55 (m, 2 H), 1.24-1.31 (m, 1 H). LCMS *m/z* calc. for $C_{29}H_{32}N_6O$ $[M+H]^+$ 481.3, found 481.4. 13C NMR (101 MHz, CHLOROFORM-d) δ ppm 14.46 (s, 1 C) 18.43 (s, 1 C) 34.00 (s, 1 C) 39.17 (s, 1 C) 46.07 (s, 1 C) 49.09 (s, 1 C) 54.92 (s, 1 C) 114.47 (s, 1 C) 117.58 (s, 1 C) 120.61 (s, 1 C) 123.89 (s, 1 C) 123.96 (s, 1 C) 125.68 (s, 1 C) 125.72 (s, 1 C) 126.19 (s, 1 C) 129.18 (s, 1 C) 129.82 (s, 1 C) 129.98 (s, 1 C) 131.59 (s, 1 C) 136.69 (s, 1 C) 138.47 (s, 1 C) 145.95 (s, 1 C) 149.05 (s, 1 C) 149.15 (s, 1 C) 152.09 (s, 1 C) 170.63 (s, 1 C).

PXRD pattern:

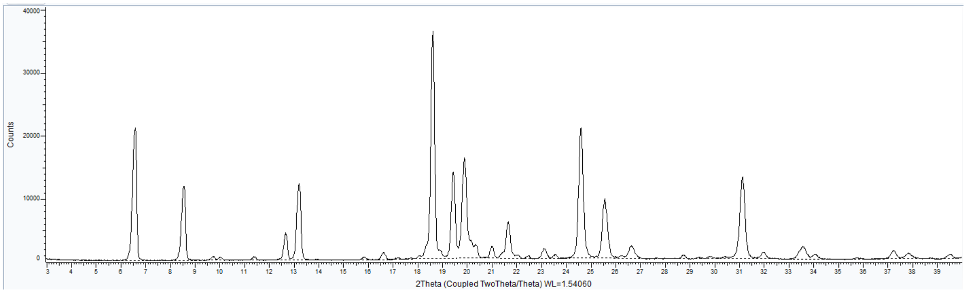

**Synthesis of compound 2 analogs (library in fig. 1B)**

Analogs of **2** were prepared using parallel chemistry employing a Palladium catalyzed Suzuki coupling condition from N-(1-(2-chloroquinolin-4-yl)ethyl)-2-methylbenzamide and boronates. The procedure is described below:

1. Dispense (N-(1-(2-chloroquinolin-4-yl)ethyl)-2-methylbenzamide (60 µmol, 1.0 eq.) and the desired boronic acid/ester Ar-B(OR)(OR) (90 µmol, 1.5 eq.) to 8 mL reaction vials.

2. Dispense 450 μL dioxane to each vial.
3. Dispense Pd-118 (6.0 μmol, 0.1 eq.) to each vial under $N_2$.
4. Dispense 150 μL $K_3PO_4$ (2.0 M, 300 μmol, 5.0 eq.) to each vial.
5. Cap vials and shake at 80 °C for 16 hours.
6. Spot check reactions by LC-MS.
7. Remove solvent by Speedvac.
8. Wash the mixture with 1.0 mL of $H_2O$ and extract with EtOAc (2.0 mL x 3).
9. Collect organic layer and evaporate solvent by Speedvac.
10. Purify residue by preparative HPLC to give desired product.

**Synthesis of compound 3 analogs (library in fig. 1C):**

Analogs of **3** were prepared in one of two methods. In some cases, the C-N coupling was performed prior to the final step amide coupling as in the preparation of compound 3 and 4. In other cases, the C-N coupling was performed in the final step using similar conditions to step 1 of the synthesis of int-1.

#### Results

| Protease | PF-07957472 IC50 (uM) |
|---|---|
| Human Cathepsin B | >100 |
| Human Chymotrypsin | >100 |
| Human Thrombin | >100 |
| Human Caspase 2 | >100 |
| Human Cathepsin D | >100 |
| Human Cathepsin L | >100 |
| Human Cathepsin S | >100 |
| Human Cathepsin V | >100 |
| Human Cathepsin K | >100 |
| Human Cathepsin F | >100 |
| Human Immunodeficiency Virus-1 | >100 |
| Human Elastase | >100 |

**Table S3.** Selectivity of PF-07957472 against mammalian and HIV proteases. The inhibitory activity of PF-07957472 was evaluated using a FRET-based assay format at several mammalian cysteine (caspase 2, cathepsin B, cathepsin L, cathepsin S, cathepsin

V, cathepsin K, cathepsin F), serine (chymotrypsin, elastase, thrombin) and aspartyl (cathepsin D, HIV-1) proteases. Data shown represent at least two independent experiments.

| Virus | PF-07957472<br>Ki (95% CI) nM |
|---|---|
| SARS-CoV-2 | 1.9 (1.1-3.2) |
| SARS-CoV-1 | 1.6 (1.3-1.9) |
| 229E-CoV | >10,000 |
| MERS-CoV | >10,000 |
| OC43-CoV | >10,000 |

**Table S4**. PF-07957472 inhibition of human coronavirus papain-like proteases.

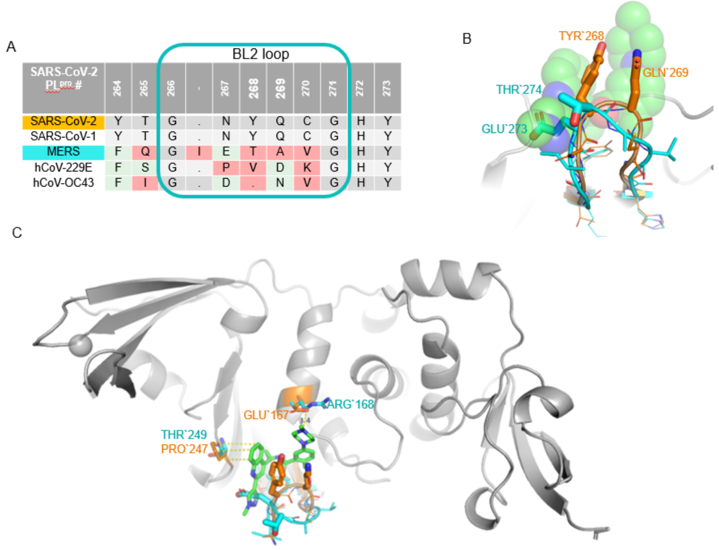

**Figure S3.** BL2 binding loop from human coronavirus papain-like proteases. (A) sequence alignment of the papain-like proteases from SARS-CoV-2, SARS-CoV-1, MERS, 229E and OC43. The non-conserved amino acids are highlighted in red boxes, green/blue indicate sharing similarity while identical residues are kept in a grey background. (B) Structural comparison of BL2 loop of $PL^{pro}$ from SARS-CoV-2 (orange pdb ID this study) and MERS (cyan, pdb ID 4RF0) showing the high flexibility in the loop and sequence variability especially betweenTyr268/Gln269 from SARS-CoV that make direct interactions with compound **4** (green) and Glu273/Thr274 from MERS. (C) Full view of $PL^{pro}$ indicating two ligand contact residues that vary between SARS-CoV-2 and MERS and are important for charge interaction (Glu167-->Arg168) and form the proline shelf (Aro247 --> Thr249)

| **Cell line** | **Mean CC50 (μM)** |
|---|---|
| Vero E6 | 69.3, n=3 |
| Vero E6 + 2 M | 78.7, n=3 |

| CP-100356 |
|---|

**Table S5**. Cytotoxicity testing of PF-07957472 in recombinant cell lines used for antiviral assays. Cytotoxicity of PF-07957472 was evaluated in non-infected Vero E6 enriched for ACE2. A P-glycoprotein inhibitor, CP-100356 was added at dose indicated to inhibit the efflux of PF-07957472 from Vero cells. Compound cytotoxicity was calculated as CC50 relative to DMSO treated and a cytotoxic control compound.

| Species[a] | Dose (mg/kg) | Dose Formulation | $C_{max}$ (ng/ml) | $T_{max}$ (h) | $AUC_{0-\infty}$ (ng·h/ml) | $CL_p$ (ml/min/kg) | Vdss (L/kg) | $t_{1/2}$ (h) | Oral F (%) |
|---|---|---|---|---|---|---|---|---|---|
| Male C57BL6 mouse | 1.0 (iv) | solution, 10% DMSO / 90% (23% w/v) HPBCD in DI water (v/v) | --- | --- | 708 (765, 650) | 23.7 (21.8, 25.6) | 2.13 (1.97, 2.28) | 3.290 (3.35, 3.22) | --- |
| | 10 (po) | solution, 2% Tween80: 98% (0.5% methylcellulose) | 862 (703, 1020) | 2.00 (2.00, 2.00) | 3760 (4300, 3220) | --- | --- | 2.19 (2.11, 2.28) | 53.10 (60.8, 45.5) |
| Male Wistar Hanover Rat | 1.0 (iv) | solution, 10% DMSO / 90% (23% w/v) HPBCD in DI water (v/v) | --- | --- | 668 (599, 737) | 25.2 (27.8, 22.6) | 4.42 (4.96, 3.88) | 3.59 (3.76, 3.41) | --- |
| | 10 (po) | solution, 2% Tween80: 98% (0.5% methylcellulose) | 435 (498, 371) | 3.00 (2.00, 4.00) | 4650 (6030, 3270) | --- | --- | 3.19 (3.16, 3.21) | 69.7 (90.3, 49.0) |
| Male Beagle Dog | 0.5 (iv) | solution, 10% PEG400/90% of 23% HPBCD | --- | --- | 640 (534, 745) | 13.4 (15.6, 11.2) | 1.59 (1,49, 1.68) | 2.06 (1.22, 2.91) | --- |
| | 3.0 (po) | suspension, 2% Tween80: 98% (0.5% methylcellulose) | 354 (323, 385) | 1.00 (1.00, 1.00) | 1580 (1280, 1870) | --- | --- | 2.52 (2.61, 2.44) | 41.0 (33.4, 48.7) |
| Male Cynomolgus Monkey | 1.0 (iv) | solution, 5% PEG400 / 95% HPBCD | --- | --- | 1600 (1840, 1350) | 10.7 (9.06, 12.3) | 2.09 (1.81, 2.38) | 3.56 (3.66, 3.45) | --- |

**Table S6**. Low-dose Preclinical pharmacokinetics of PF-07957472
[a] Pharmacokinetic parameters were calculated from plasma concentration–time data and are reported as mean (± S.D. for n=3 and individual values for n=2). All pharmacokinetics were conducted in male sex of each species. Oral (po) studies were conducted in the fed state unless otherwise noted.

|  | PF-07957472 Mouse Multiple Doses |  |  |  |  |  |
|---|---|---|---|---|---|---|
| Route | IV | PO | PO | PO | PO | PO |
| Dose (mg/kg) | 1 | 10 | 15 | 30 | 50 | 150 |
| Dose Vehicle | 10% DMSO / 90% (23% w/v) HPBCD in DI water (v/v) | 2% Tween80: 98% (0.5% methylcellulose) | 2% (v/v) Tween80 in 98% (v/v) of 0.5% (w/v) MethylCellulose | 2% Tween80: 98% (0.5% methylcellulose) | 2% (v/v) Tween80 in 98% (v/v) of 0.5% (w/v) MethylCellulose | 2% (v/v) Tween80 in 98% (v/v) of 0.5% (w/v) MethylCellulose |
| mouse strain | C57BL6 | C57BL6 | Balb c | C57BL6 | Balb c | Balb c |
| mouse sex | Male | Male | Female | Male | Female | Female |
| Dose Formulation | Solution | Solution | Suspension | Solution | Suspension | Suspension |
| AUC(0-Tlast) (ng*Hours/mL) | 706 | 3760 | 7940 | 14300 | 29700 | 89100 |
| AUC(0,inf) (ng*Hours/mL) | 708 | 3760 | 7940 | 14300 | 29800 | 89200 |
| Cmax (ng/mL) | - | 862 | 1140 | 1880 | 5410 | 12800 |
| Tmax (Hours) | - | 2 | 1.25 | 2 | 0.375 | 0.25 |
| T1/2 (Hours) | 3.29 | 2.19 | 1.95 | 2.17 | 2.01 | 2.47 |
| Vdss (L/kg) | 2.13 | - | - | - | - | - |
| CL minutes (mL/min/kg) | 23.7 | - | - | - | - | - |
| % F Method 3 (Dose * AUC Extrap) | - | 53.1 | 74.8 | 67.4 | 84.0 | 84.0 |
| % of dose in feces | 26.0% | 12.6% | - | 10.8% | - | - |

**Table S7: Mouse multi-dose Formulation**

Oral pharmacokinetic studies were conducted in the fed state unless otherwise noted. Oral rat pharmacokinetic studies were conducted with xx or xx form xx. The aforementioned low-dose 1 mg/kg IV was formulated as solutions in 10% DMSO / 90% (23% w/v) HPBCD in DI water (v/v). The 10 mg/kg PO doses are included for comparison. The For the 15, 50, 150 mg/kg doses, a homogeneous suspension formulation in 2% (v/v) Tween80 in 98% (v/v) of 0.5% (w/v) MethylCellulose by utilizing Geometric dilution were prepared.

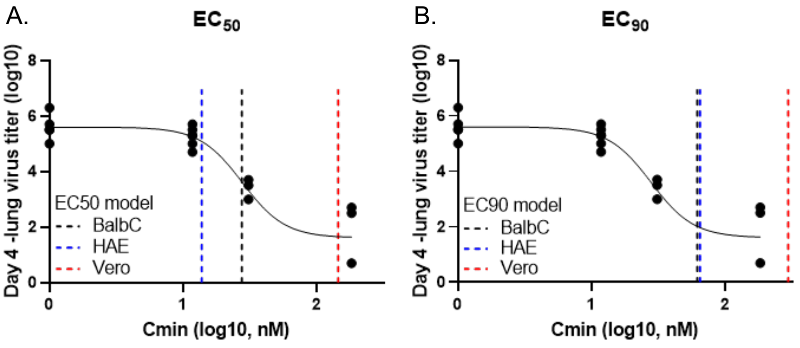

C.

|  | EC50 (nM) | EC90 (nM) |
|---|---|---|
| BalbC | 28 | 62 |
| HAE | 14 | 65 |
| Vero | 147 | 296 |

**Figure S4 -EC50. In vitro – in vivo – correlation of compound potency.** The $EC_{50}$ (A) and $EC_{90}$ (B) were calculated for the compound in BalbC mice based on the projected Cmin and lung virus titer after 4 days of treatment (points = data, solid black line = fitted sigmoidal curve). The EC50 and $EC_{90}$s calculated from Vero E6 (red) and HAE (blue) cells were overlayed and revealed that the potency measured in HAEs and mice were well-aligned, but the potency measured in Vero E6 cells was right shifted. (C) Relative to in BalbC mice, the $EC_{50}$ and $EC_{90}$ from HAEs were within 2fold, while those from Vero E6 ~5fold larger. These results supported driving the Cmin threshold off of HAE $EC_{90}$ instead of Vero E6.

**Table S8. Mutation count/ frequency of SARS-CoV-2 PLPro for key contact residues within 5A around PF-07957472 and the catalytic /tunnel residues.**

| PLpro coordinate | Residue | Mutations* | Counts* | Frequency* |
|---|---|---|---|---|
| 856 | C | V, S, F, L, R, Y, * | 23 | 3.38E-06 |
| 857 | Y | H, C, Q, F, *, N | 117 | 1.72E-05 |
| 902 | K | N, R, *, M, I, T, E, G | 1640 | 2.41E-04 |
| 907 | L | F, S, V, *, R | 109 | 1.60E-05 |
| 908 | G | S, C, A, R, *, D, V | 216 | 3.17E-05 |
| 909 | D | A, G, E, Q, Y, N, del | 85 | 1.25E-05 |
| 910 | V | I, A, F, G, P, D, M, S | 351 | 5.16E-05 |
| 911 | R | K, L, G, S | 53 | 7.79E-06 |
| 912 | E | D, G, A, K, H, R, Q, V, del | 151 | 2.22E-05 |
| 916 | Y | H, C, F, N, L, S, del | 630 | 9.26E-05 |
| 953 | M | T, I, V, L, R, K, P | 1491 | 2.19E-04 |
| 991 | A | V, T, S, E, L, N, Q, G, P | 3138 | 4.61E-04 |
| 992 | P | L, S, T, Q, I, del | 2576 | 3.78E-04 |
| 993 | P | F, L, S, H, *, R, T, Y | 206 | 3.03E-05 |
| 1009 | Y | H, P, *, F, K, C, D, del | 38 | 5.58E-06 |
| 1010 | T | I, N, A, S, P, del | 1159 | 1.70E-04 |
| 1011 | G | V, S, A, C, Y | 22 | 3.23E-06 |
| 1012 | N | S, D, Y, K, T, I, del, C | 458 | 6.73E-05 |
| 1013 | Y | H, C, del, R, N, D, S, F, * | 1318 | 1.94E-04 |
| 1014 | Q | R, L, K, E, H, V, * | 169 | 2.48E-05 |
| 1015 | C | Y, R, F, S, G, W | 527 | 7.74E-05 |
| 1016 | G | N, S, T, C, D, V | 11 | 1.62E-06 |
| 1017 | H | N, L, Q | 7 | 1.03E-06 |
| 1018 | Y | F, *, A, S, H, Q, T, V | 19 | 2.79E-06 |
| 1046 | T | A, L, del, M, E, Q, R, S | 129 | 1.90E-05 |
| 1047 | D | N, E, *, Q, G, del, R, Y | 198 | 2.91E-05 |

* The mutation annotation was performed on 6,806,275 Omicron genome sequences collected from GISAID as of 12/08/2023

**Table S9. Media binding of PF-07957472 for in vitro cell systems (n = 4)**

| | Fu (CV) | media |
|---|---|---|
| PF-07957472 | 0.749 (10%) | 2% HI-FBS in DMEMEpiAirway |
| PF-07957472 | 0.916 (10%) | HAE maintenance media from Mattek |

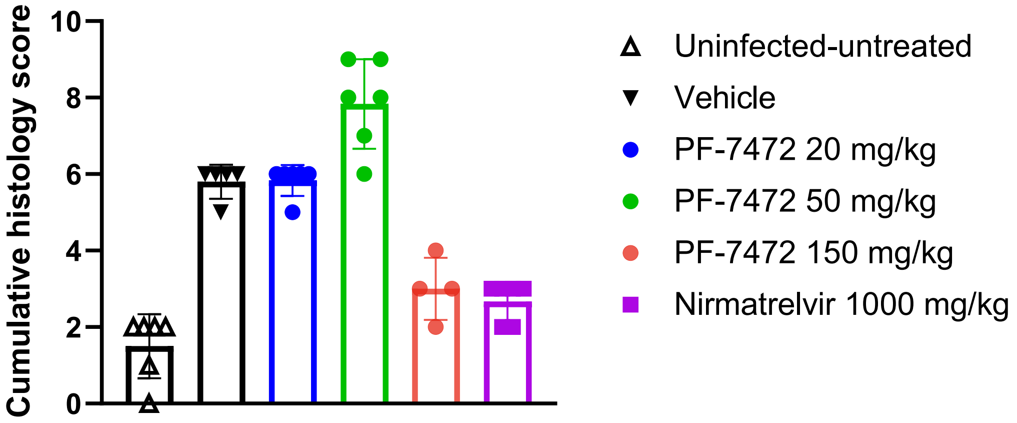

**Figure S5.** Lung Histopathology score from lung histopathology of mouse-adapted SARS-CoV-2 infection 4 days post dose of PF-07957472 at 20, 50, and 150 mg/kg, Nirmatrelvir at 1000 mg/kg, vehicle, or uninfected, untreated animals. Histopathology score parameters include perivascular inflammation, bronchiolar epithelial degeneration or necrosis, bronchiolar inflammation, and alveolar inflammation.

1 Ghosh, A. K. *et al.* Structure-Based Design, Synthesis, and Biological Evaluation of a Series of Novel and Reversible Inhibitors for the Severe Acute Respiratory Syndrome−Coronavirus Papain-Like Protease. *J. Med. Chem.* **52**, 5228-5240 (2009).
2 Owen, D. R. *et al.* An oral SARS-CoV-2 M(pro) inhibitor clinical candidate for the treatment of COVID-19. *Science* **374**, 1586-1593 (2021).
3 Natekar, J. P. *et al.* Differential Pathogenesis of SARS-CoV-2 Variants of Concern in Human ACE2-Expressing Mice. *Viruses* **14** (2022).
4 Maurer, T. S., Smith, D., Beaumont, K. & Di, L. Dose Predictions for Drug Design. *J. Med. Chem.* **63**, 6423-6435 (2020).
5 Doran, A. C. *et al.* Defining the Selectivity of Chemical Inhibitors Used for Cytochrome P450 Reaction Phenotyping: Overcoming Selectivity Limitations with a Six-Parameter Inhibition Curve-Fitting Approach. *Drug Metab. Disp.*, DMD-AR-2022-000884 (2022).
6 Leist, S. R. *et al.* A Mouse-Adapted SARS-CoV-2 Induces Acute Lung Injury and Mortality in Standard Laboratory Mice. *Cell* **183**, 1070-1085.e1012 (2020).
7 REED, L. J. & MUENCH, H. A SIMPLE METHOD OF ESTIMATING FIFTY PER CENT ENDPOINTS12. *American Journal of Epidemiology* **27**, 493-497 (1938).
8 Freitas, B. T. *et al.* Characterization and Noncovalent Inhibition of the Deubiquitinase and deISGylase Activity of SARS-CoV-2 Papain-Like Protease. *ACS Infectious Diseases* **6**, 2099-2109 (2020).
9 Lee, H. *et al.* Inhibitor Recognition Specificity of MERS-CoV Papain-like Protease May Differ from That of SARS-CoV. *ACS Chemical Biology* **10**, 1456-1465 (2015).
10 Ratia, K. *et al.* A noncovalent class of papain-like protease/deubiquitinase inhibitors blocks SARS virus replication. *Proc Natl Acad Sci U S A* **105**, 16119-16124 (2008).
11 Ziebuhr, J. *et al.* Human coronavirus 229E papain-like proteases have overlapping specificities but distinct functions in viral replication. *J Virol* **81**, 3922-3932 (2007).
12 Lei, J., Kusov, Y. & Hilgenfeld, R. Nsp3 of coronaviruses: Structures and functions of a large multi-domain protein. *Antiviral Res* **149**, 58-74 (2018).
13 Fu, Z. *et al.* The complex structure of GRL0617 and SARS-CoV-2 PLpro reveals a hot spot for antiviral drug discovery. *Nature Communications* **12**, 488 (2021).
14 Vonrhein, C. *et al.* Data processing and analysis with the autoPROC toolbox. *Acta Crystallogr D Biol Crystallogr* **67**, 293-302 (2011).

15 McCoy, A. J. *et al.* Phaser crystallographic software. *J. Appl. Crystallogr.* **40**, 658-674 (2007).
16 Emsley, P., Lohkamp, B., Scott, W. G. & Cowtan, K. Features and development of Coot. *Acta Crystallogr D Biol Crystallogr* **66**, 486-501 (2010).
